# Protistan grazing impacts microbial communities and carbon cycling at deep-sea hydrothermal vents

**DOI:** 10.1101/2021.02.08.430233

**Authors:** Sarah K. Hu, Erica L. Herrera, Amy R. Smith, Maria G. Pachiadaki, Virginia P. Edgcomb, Sean P. Sylva, Eric W. Chan, Jeffrey S. Seewald, Christopher R. German, Julie A. Huber

## Abstract

Microbial eukaryotes (or protists) in marine ecosystems are a link between microbial primary producers and all higher trophic levels. The rate at which heterotrophic protistan grazers consume microbial prey and recycle organic matter is an important factor that influences marine microbial food webs and carbon cycling. At deep-sea hydrothermal vents, chemosynthetic bacteria and archaea form the base of a food web that functions in the absence of sunlight, but the role of protistan grazers in these highly productive ecosystems is largely unexplored. Here, we pair grazing experiments with a molecular survey to quantify protistan grazing and to characterize the composition of vent-associated protists in low-temperature venting fluids from Gorda Ridge in the North East (NE) Pacific Ocean. Results reveal protists exert higher predation pressure at vents compared to the surrounding deep seawater environment and may account for consuming 28-62% of the daily stock of prokaryotic biomass within the hydrothermal vent food web. The vent-associated protistan community was more species rich relative to the background deep sea, and patterns in the distribution and co-occurrence of vent microbes provide additional insights into potential predator-prey interactions. Ciliates, followed by dinoflagellates, Syndiniales, rhizaria, and stramenopiles dominated the vent protist community and included bacterivorous species, species known to host symbionts, and parasites. Our findings provide an estimate of protistan grazing pressure within hydrothermal vent food webs, highlighting the role that diverse deep-sea protistan communities have in carbon cycling.

**Significance:** Heterotrophic protists are ubiquitous in all aquatic ecosystems and represent an important ecological link because they transfer organic carbon from primary producers to higher trophic levels. Here, we quantify the predator-prey trophic interaction among protistan grazers and microbial prey at multiple sites of hydrothermal venting near the Gorda Ridge spreading center in the NE Pacific Ocean. Grazing pressure was higher at the site of active diffuse flow and was carried out by a highly diverse assemblage of protistan species; elevated grazing rates are attributed to higher concentrations of chemosynthetic microorganisms and biological diversity localized to hydrothermal vent environments.

## Introduction

Mixing of venting hydrothermal fluids with surrounding seawater in the deep sea creates redox gradients that promote a hub of biological activity supported by chemosynthetic primary production in the absence of sunlight. These localized regions of elevated microbial biomass are important sources of carbon and energy to the surrounding deep-sea ecosystem (1–5). In particular, the consumption of hydrothermal vent microorganisms by single-celled microbial eukaryotes (or protists) is an important link in the food web where carbon is transferred to higher trophic levels or remineralized to the microbial loop.

Protistan grazing is a significant source of mortality for bacterial and archaeal populations in aquatic ecosystems that also influences their composition and diversity (6). Assessments of grazing in the mesopelagic and dark ocean have found that rates of consumption decrease with depth and correspond to bacterial abundance (7, 8). Therefore, at sites of increased biological activity and microbial biomass, such as areas of redox stratification, protistan grazing is higher relative to the rest of the water column (9, 10). Comparable data have been lacking from deep-sea hydrothermal vents, where the relatively high microbial biomass and rates of primary productivity suggest protistan grazing should be a significant source of microbial mortality and carbon transfer. Further, single-celled microbial eukaryotes can serve as a nutritional resource for other larger protists and higher trophic levels (4, 11).

Early microscopic and culture-based experiments from several hydrothermal vents confirmed the presence of single-celled microbial eukaryotes, with observations and enrichment cultures revealing diverse assemblages of flagellated protists and ciliates (12, 13). The study of protistan taxonomy and distribution via genetic analyses at deep-sea vents has uncovered a community largely composed of alveolates, stramenopiles, and rhizaria (14–17). In addition to many of these sequence surveys identifying known bacterivorous species, ciliates isolated from Guaymas Basin were shown to consume an introduced prey analog (18). Collectively, these studies provide supporting evidence of a diverse community of active protistan grazers at deep-sea vents.

Here, we investigate protistan predation pressure upon microbial populations in venting fluids along the Gorda Ridge to test the hypothesis that protistan grazing and diversity is elevated within hydrothermal habitats compared to the surrounding deep sea due to increased prey availability. Estimates of mortality via protistan phagotrophy are calculated from grazing experiments conducted with low temperature diffusely venting fluid that mixes with seawater at and below the seafloor. Paired 18S rRNA gene amplicon sequencing from the same experimental sites and incubations reveal the *in situ* protistan diversity and distribution to evaluate potential preferences in prey, with a focus on the protistan grazer population and their relationship to bacteria and archaea. We present quantitative estimates of protistan grazing from a deep-sea hydrothermal vent ecosystem, thus providing new details into the role protists play in food webs and carbon cycling in the deep sea.

## Results & Discussion

### Sea Cliff and Apollo hydrothermal vent fields

Low-temperature (10-80°C) diffusely venting fluids were collected at the Sea Cliff and Apollo hydrothermal vent fields along the Gorda Ridge (Figure S1; 19, 20, 21). Hydrothermal vent fluids collected for experiments and genetic analysis were geochemically distinct from plume (5 m above active venting), near vent bottom water (lateral to venting fluid), and background seawater (outside the range of hydrothermal influence; Table 1). The concentration of bacteria and archaea was 5-10 × 10^4^ cells ml^-1^ in low temperature vent fluids, which was higher than background seawater concentrations (3-5 × 10^4^ cells ml^-1^; Table 1). Diffuse vents sampled in both fields represented a mixture of nearby high temperature vent fluid (Candelabra, 298°C and Sir Ventsalot, 292°C) with seawater (22). During sample collection (30-40 minutes), the temperature of the fluid being sampled fluctuated between 3-72°C, due to mixing (Table 1). The temperature maxima at Mt. Edwards and Venti Latte were lower compared to Candelabra and Sir Ventsalot, and ranged from 11-40°C; these sites also had visible tube worm clusters (*Paralvinella palmiformis*; Figure S1; Table 1).

**Table 1.** Chemical characteristics of the samples used in this study, as well as nearby high-temperature end members. For vents sampled with both the SUPR and IGT, data in [brackets] are from SUPR bag samples used in grazing experiments, whereas all other data is from paired IGT samples at the same site. Sir Ventsalot was only sampled via SUPR. For plume and seawater samples, data are from Niskin bottles.

| Vent Site | Depth | Tmax<br>[range] | pH | Mg | %<br>Seawater<br>(bag<br>sample) | H<sub>2</sub>S | H<sub>2</sub> | CH<sub>4</sub> | Microbial |
| --- | --- | --- | --- | --- | --- | --- | --- | --- | --- |
|  | (m) | (°C) |  | (mM) |  | (mM) | (uM) | (uM) | (cell mL<sup>-1</sup>) |
| Candelabra High Temperature Vent, Gorda Vent Field | 2730 | 298 | 4.5 | 2.1 | --- | 3 | 62.0 | 68.4 | --- |
| Candelabra Diffuse Vent | 2730 | 79 [9-68] | 5.5 [5.8] | 35.7 [45.8] | 88.4% | n.d. | 21.9 | 23.7 | 5.51E+04 |
| Candelabra Plume | 2725 | 1.7 | --- | --- | --- | --- | --- | --- | 7.69E+04 |
| Venti Latte Vent | 2708 | 11 [10-23] | 6.4 [5.5] | 50.9 [50.4] | 97.3% | n.d. | b.d. | 0.9 | 1.11E+05 |
| Mt Edwards Vent | 2707 | 40 [15-30] | 6.0 [5.8] | 42.6 [42.8] | 82.5% | 1.01 | 127.0 | 10.1 | 5.14E+04 |
| Mt Edwards Plume | 2702 | 1.8 | --- | --- | --- | --- | --- | --- | --- |
| Sir Ventsalot High Temperature Vent, Apollo Vent Field | 2732 | 292 | 2.8 | 2.5 | --- | 2.53 | 71.4 | 66.7 | --- |
| Sir Ventsalot Diffuse Vent | 2732 | [3-72] | n.d. | 50.8 | 98.0% | --- | --- | --- | 5.30E+04 |
| Near vent bottom water | 2745 | 1.7 | 7.8 | 51.8 | 100% | --- | --- | --- | 5.20E+04 |
| Seawater: Shallow | 150 | 8.6 | n.d. | 51.8 | 100% | --- | --- | --- | --- |
| Seawater: Deep | 2010-2090 | 1.8 | 7.8 | 51.8 | 100% | --- | --- | --- | 3.91E+04 |

*n.d., no data; b.d., below detection*

### Protistan grazers exert predation pressure on hydrothermal vent bacteria and archaea

Grazing incubations conducted with fluids collected from five sites within the Sea Cliff and Apollo vent fields demonstrate that microbial eukaryotes actively graze microbial communities in hydrothermal vent fluids at an elevated rate relative to the surrounding deep-sea environment (Figure 1). Protists consumed microbial prey at rates ranging between 700 to 1828 cells ml^-1^ hour^-1^ in the diffuse venting fluids (Figure 1b, Table S1), whereas in near vent bottom water away from active venting, the grazing rate was 255 cells ml^-1^ hour^-1^. The prokaryote turnover rate, expressed as the percentage of the daily consumed prokaryotes relative to the standing stock (average prokaryotic concentration), was 17.2% in the bottom water near the hydrothermal vent sites. Protistan grazing at hydrothermal vents accounts for 28-62% of the daily prokaryote biomass turnover (Figure 1c; Table S1), demonstrating that the vent microbial community is under more top-down pressure compared to communities in the background deep-sea environment.

**Figure 1.**
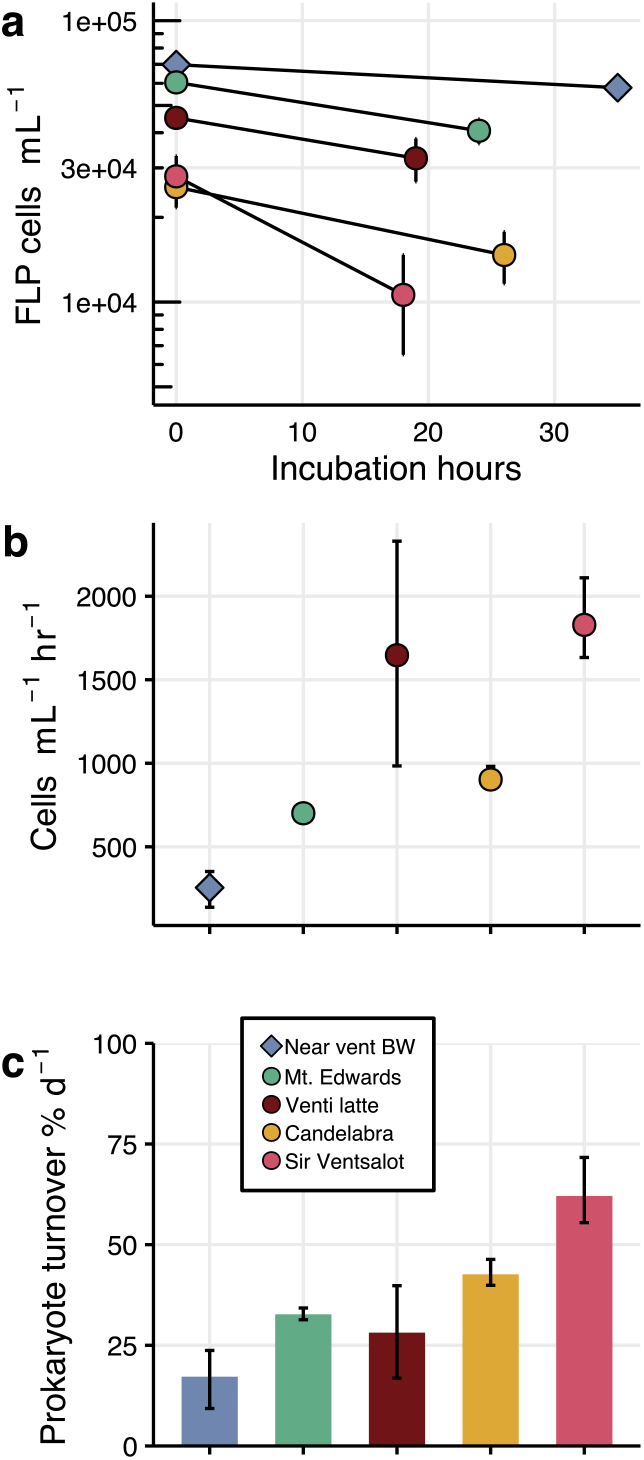
Results from grazing experiments conducted at Sea Cliff and Apollo hydrothermal vent fields. (**a**) Loss of Fluorescently-Labelled Prey (log FLP cells ml^-1^; y-axis) during each incubation (hours; x-axis). Error bars represent the standard mean error from the average across replicates. (**b**) Grazing rate for each site expressed as the consumption of cells ml^-1^ hr^-1^, derived from Equation (1). Error bars report the minimum and maximum grazing rate derived from the standard mean error. (**c**) Estimated daily prokaryote turnover percentage (% d^-1^), where the grazing rate for each site was multiplied by the *in situ* prokaryote cell concentration (Table 1). Error bars represent the minimum and maximum derived from the range of grazing rate for each incubation. Complete experiment details are reported in Table S1.

Free-living heterotrophic protists may adapt to low prey encounter rates, due to decreased microbial biomass in the deep sea, by associating with sinking particles or localized habitats with more abundant prey (6, 23). Transition zones such as redoxclines are also sites of comparatively higher grazing pressure since they host a more diverse and abundant prokaryotic community due to the presence of diverse sources of carbon and energy (8, 10, 24). In one of the only other studies to quantify deep-sea predation pressure, protistan grazing within a deep-sea halocline (3500 m; above the hypersaline Urania Basin in the Eastern Mediterranean Sea) was calculated to be over 13,500 cells ml^-1^ hour^-1^, in contrast to 10-390 cells ml^-1^ hour^-1^ in the water column outside the influence of the halocline (100-3000 m; 9). Near vent bottom water grazing rates in our study (Figure 1; Table S1) were comparable to rates previously obtained from mesopelagic and bathypelagic water column depths (200-2500 m; ∼10-400 cells consumed ml^-1^ hour^-1^; (8, 10)), while grazing pressure and prokaryotic abundance were higher in vent fluids (Figure 1; Table S1).

Commensurate with typical declining concentrations of prokaryotes with ocean depth, deep-sea grazing rates in this study were lower relative to those measured in seawater from euphotic regions (10, 25). However, when the microbial community biomass is taken into account, the impact of protistan grazing measured as a daily prokaryote turnover rate (28-62% day^-1^) at the hydrothermal vent sites are within the range of turnover rates reported from some euphotic zone studies (6). This observation is consistent with grazing rates reported from sub-euphotic depths, especially at environments with increased biological activity (reviewed in 10). Our results demonstrate that protistan grazing plays an important role in processing new organic carbon within hydrothermal vent food webs. Furthermore, daily prokaryote turnover rates observed in this study correlate positively with temperature maxima at each vent site (r^2^ = 0.87; Figure S4b). While additional work is required to comprehensively link protistan grazing activity to hydrothermal vent geochemistry, results from previous studies also found protistan grazing to positively correspond with temperature (25, 26).

Grazing rates from Sea Cliff and Apollo vent fields indicate that protists may be consuming or remineralizing 1.45 - 3.77 µg of carbon L^-1^ day^-1^ (Table S1; using a carbon conversion factor of 86 fg carbon cell^-1^ (27)). While few measurements of absolute fixed carbon exist from hydrothermal vents, McNichol *et al*. estimate primary production by the fluid-associated microbial community within low temperature diffuse fluids at the East Pacific Rise to range between 17.3 - 321.4 µg C L^-1^ day^-1^, at 24°C and 50°C under *in situ* pressure, representing an important source of new labile carbon in the deep sea (2, 3). Considering these estimates, protistan grazing may account for consumption, or transformation of up to 22% of carbon fixed by the chemosynthetic population. The eventual fate of this carbon contributes to the organic carbon flux that reaches the seafloor; locally, the flux may outweigh the supply of sinking organic carbon from the surface ocean (1). Our findings show that the link between microbial prey abundance and activities of protistan grazers at hydrothermal vents is significant and may account for a substantial amount of organic carbon transfer at the base of deep-sea food webs.

### Distinct microbial populations at hydrothermal vents

The Sea Cliff and Apollo hydrothermal vent sites were found to host a diverse assemblage of protists (Figures 2a and 3). Amplicon sequencing of the protistan (18S rRNA gene) and prokaryotic (16S rRNA gene) communities resulted in 9027 and 6497 amplicon sequence variants (ASVs), respectively. ASVs represent approximately species-level designations based on recovered sequences (see *Supplementary Text*). The taxonomic composition of 18S rRNA gene-derived ASVs reveal dominant members of the vent ecosystem to include ciliates, dinoflagellates, Syndiniales, rhizaria, and stramenopiles (Figure 2a); these same protistan groups are enriched in other deep-sea niche habitats, such as methane seeps and other hydrothermal vent systems or vent-fluid influenced environments (14, 15, 17, 18, 28). Community-wide analyses of both protists (18S rRNA gene amplicons) and bacteria and archaea (16S rRNA gene amplicons) showed that replicate samples cluster together (Figures 2b, S5, S6; *Supplementary Text*). Background, plume, and near vent bottom water bacteria and archaea community compositions were distinct from the vent-associated community (Figure S6). Sites of actively venting fluid hosted higher relative sequence abundances assigned to the *Epsilonbacteraeota* class, including *Sulfurimonas* and *Sulfurovum* (Figure S6a), which are commonly dominant within vent microbial communities (29). The expected impact of vent fluid collection and depressurization was evidenced by differences in the protistan community composition in samples from *in situ* (SUPR or sterivex filters) and the start of each grazing experiment (T_0_; Figure 2a) (30). However, consistency among grazing experiment sample community composition and ordination analysis demonstrated that the collected vent fluid used for grazing incubations was representative of a hydrothermally-influenced community (Figures 2a and 2b).

**Figure 2.**
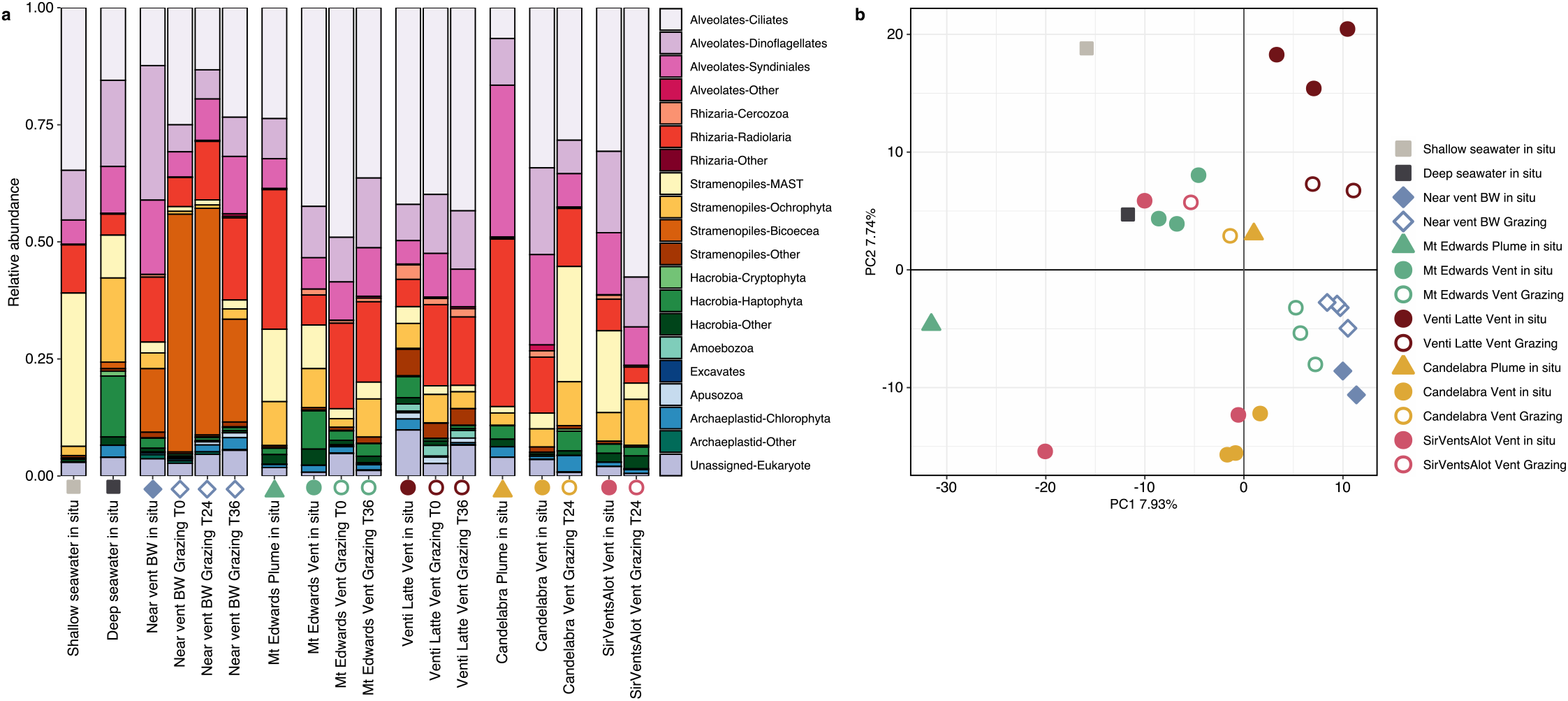
Summary of protistan diversity for *in situ* and grazing experiment samples. (**a**) Taxonomic breakdown of samples, including background, plume (5 m above active flow), near vent bottom water (BW), *in situ* vent sites, and associated grazing incubation bottles (T_x_). Bar plot reports the relative sequence abundances, where colors designate major protistan taxonomic groups (based on manual curation, see *Materials and Methods*). (**b**) Ordination analysis of all samples, including replicates, from the 18S rRNA gene-derived sequence data. Data was center log-ratio transformed ahead of PCA analysis. For both (**a**) and (**b**), symbols indicate origin of sample and color denotes vent site. Solid symbols represent *in situ* samples and open symbols designate samples from grazing experiments. Samples from grazing experiments include different time points (Table S1).

To test the hypothesis that microbial eukaryotes from the surrounding deep-sea environment have greater species richness at sites of low temperature diffuse venting fluid, ASVs were classified based on their distribution among vent fluid and non-hydrothermally influenced environments (background). ‘Resident’ ASVs were found only within hydrothermally-influenced samples and considered to be potentially vent endemic, and ‘cosmopolitan’ ASVs included those detected throughout the background and hydrothermally-influenced samples (*Supplementary Text*; Figure S7). The total number of ASVs within the resident population was several fold higher than in the cosmopolitan population (4236 resident versus 535 cosmopolitan ASVs), yet the number of sequences within each population was similar (48% cosmopolitan and 46% resident). This trend was also observed in 18S rRNA gene surveys from Mariana Arc vent fluids (17). While biases with sequence-based analyses inhibit our ability to infer absolute abundances and do not necessarily provide full coverage of the entire microbial community, these results suggest that putative vent endemic and cosmopolitan taxa have similar abundances within the protistan hydrothermal vent community, but the resident protistan population is comparatively more species diverse (Figure 3). Combined with the elevated grazing rates within vent fluid, results from the molecular survey support our hypothesis that protists are enriched in hydrothermal mixing zones relative to the surrounding deep-sea environment due to increased prey availability.

**Figure 3.**
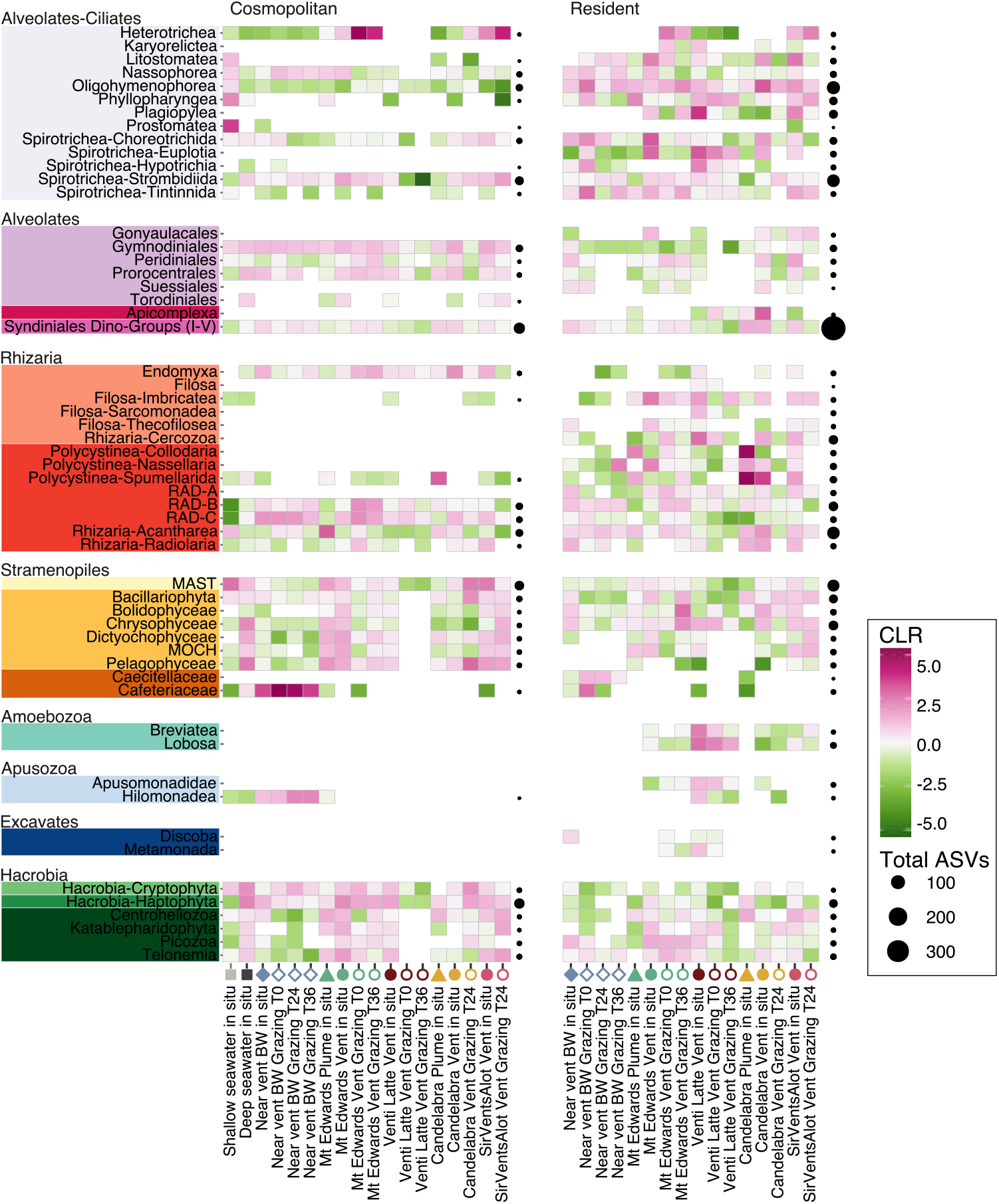
Prevalence, distribution, and richness of protists at the class or order level across all samples. Centered log-ratio (CLR) transformed sequence abundances (green to pink heat map) for all samples, including background, *in situ*, plume, and grazing incubations (columns; x-axis), by taxonomic group (color schema by row; y-axis), and classification as either cosmopolitan (left) or resident (right). CLR value is a result of transforming the sequence counts so the geometric mean equates to zero. Ahead of sequence transformation, sequences within an ASV were averaged across replicates, then sequences were summed at approximately the class or order level (y-axis). Blank spaces indicate that no sequences were detected. Bubble plots to the right of each panel represent the total number of ASVs (by size) for the distribution (cosmopolitan versus resident) for each row.

To assess the composition of the putative grazer population, we closely examined the diversity and distribution of key protistan lineages known to exhibit heterotrophy in other environments (see *Supplementary Text* for detailed observations by taxonomic group; Tables S3-S5). Ciliates were identified as important grazers in the hydrothermal vent fluids from these sites, as many groups detected include well-known bacterivorous species (31). The Oligohymenophorea and Spirotrichea classes were particularly enriched within Gorda vent fluids (Figures 2, 3), and species within these groups may be specially suited to thrive within the vent environment. For example, scuticociliates (a subclass within Oligohymenophorea; Table S5) have been found previously near hydrothermal vent sites (28, 32), and in addition to their heterotrophic capabilities, are known to be parasitic or to host endosymbionts (31). Ciliates found only within the vent fluid samples, such as Karyorelictea, Plagiopylea, and *Euplotia*, include species capable of thriving in low oxygen to suboxic environments with modified mitochondria (hydrogenosomes). Members of these classes are known to form close associations with methanogens or bacteria (33, 34). Taxonomic groups within ciliates and other alveolates, rhizaria (radiolaria and cercozoa), amoebozoa, apusozoa, and excavates that were detected primarily in the resident population include species that are candidates in future efforts to understand the functional traits among hydrothermal vent endemic protists; many of these same groups have previously been identified as including vent endemic species (17). Heterotrophic nanoflagellate members of the stramenopile supergroup were overwhelmingly MArine STramenopiles (MAST, in cosmopolitan and resident populations) or *Cafeteriaceae* (primarily in the near vent bottom water samples) (Figure 3); both are recognized as important bacterivores with a global distribution and often found in mesopelagic and deep sea surveys (35–37). MAST have also been found at higher relative sequence abundances within the Mariana Arc vent ecosystem and hydrothermally influenced water masses within Okinawa Trough (17, 28).

We also found evidence for parasitic populations of protists that may represent a source of mortality to the protists themselves and other small eukaryotes (i.e., metazoa) at the Gorda Ridge sites. Parasitic protists have been found to account for a significant portion of the globally distributed heterotrophic protist community (38), where the most abundant genetic signatures were affiliated with Syndiniales (also marine alveolate; MALV group). Syndiniales have been recognized as a major source of mortality for many microbial eukaryotes as well as metazoa and are typically found in association with ciliates, dinoflagellates, and rhizaria (39); our data suggests they may also represent a source of mortality among the vent protistan population (Dino Groups I-V; Figure 3). The prevalence of Syndiniales, along with other protistan lineages known to include parasitic species (e.g., ciliates, amoebozoa, or cercozoa), supports previous observations that parasitism is widespread and likely contributes carbon turnover in deep-sea food webs (40). Parasitism and grazing by microbial eukaryotes, along with other modes of microbial mortality such as viral lysis, should be included in future studies of deep-sea food web ecology.

Prey preferences and varied feeding strategies among protistan grazers can place selective pressure on the prey species composition based on morphology or other traits that impact prey nutritional value (41). Studies have shown instances of protistan grazers preferentially consuming larger cells or specific subdivisions of bacteria and archaea, like proteobacteria (42, 43). Outcomes from these selective pressures from bacterivory may shift the overall microbial community morphology towards large aggregates or filaments (44), and results from one experiment suggested that grazing could inhibit nitrification due to preferential consumption of larger nitrifying bacteria (45). Network analysis (46), using a subset of the 18S and 16S rRNA gene-derived ASVs, does not confirm exact preferred prey preferences among vent protists, but result allow speculation that putative prey among the ciliate grazer population includes the most abundant prokaryotic groups (Figure 4), i.e. *Alphaproteobacteria, Gammaproteobacteria, Nitrososphaeria*, and *Sulfurimonas*. To fully capture predator-prey relationships and the impact of grazers on hydrothermal vent species evenness, analysis of morphology and carbon content of preferred protistan prey are required.

**Figure 4.**
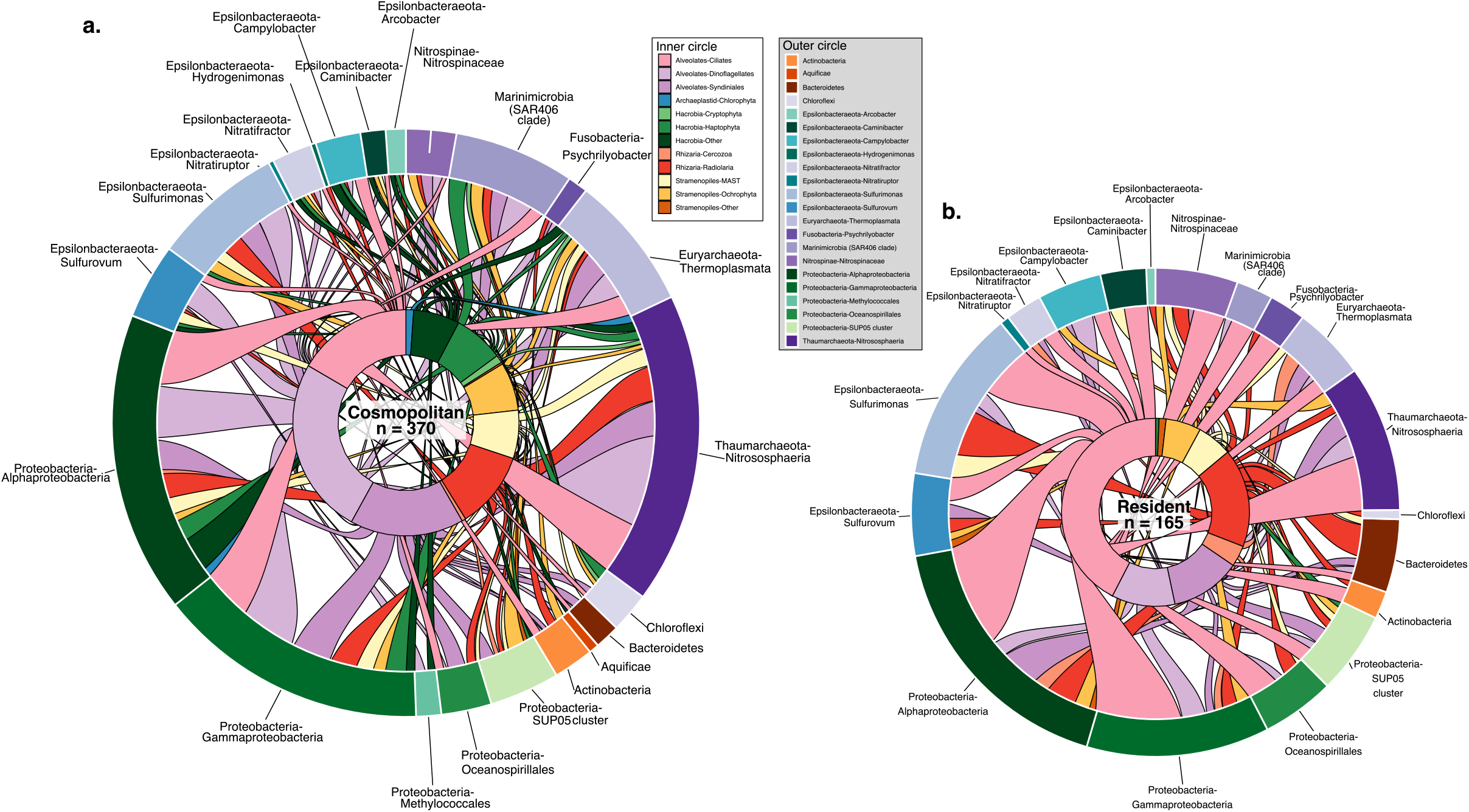
Alluvial representation of the interaction between protists (inner circle) and bacteria or archaea (outer circle) derived from the SPIEC-EASI network analysis (see *Materials and Methods*). The total number of interactions for the (**a**) cosmopolitan protistan population (n = 370) was greater than the number of interactions involving the (**b**) resident protistan population (n = 167). Color of the inner circle and alluvials that connect the inner to outer circle designate the protistan taxonomic group (derived from 18S rRNA ASVs) and the color of the outer circle represents the bacteria or archaea group (derived from the 16S rRNA ASVs). Significant 18S-16S ASV pairs are listed in Table S6, here those ASV pairs were summed together based on membership to the protistan or prokaryotic taxonomic groups.

Reported grazing rates in this study constrain the trophic exchange of protistan grazing on microbial prey at hydrothermal vents. Findings from our paired quantitative and qualitative approach provide new insight into the ecological roles of protists at deep-sea vents and their subsequent impact on the deep-sea carbon budget through carbon trophic transfer and release of dissolved organic matter. Phagotrophic grazing on smaller microorganisms accounts for a considerable amount of mortality in many aquatic environments and undoubtedly influences the diversity and composition of the hydrothermal vent microbial community; thus, efforts to fully characterize the microbial loop in the deep sea should include the roles of microbial eukaryotes. Protistan grazing is a key route of carbon transformation and exchange in the hydrothermal vent food web, and these findings contribute to our growing understanding of carbon cycling in the deep ocean.

## Materials and Methods

### Sample collection and processing

The Gorda Ridge spreading center, located ∼200 km off the coast of southern Oregon was visited in May-June 2019 with the E/V *Nautilus* (cruise NA108; 20). Low temperature diffuse hydrothermal vent fluid samples <100°C were collected using the ROV *Hercules* and a SUspended Particle Rosette Sampler (SUPR; 47). This involved measuring the fluid temperature with the *Hercules* temperature probe in regions of hydrothermal fluid flow, then positioning the sampler intake into the vent for fluid collection. The SUPR sampler pumped fluid to either fill gas-tight bags (PET/METPET/LLDPE; ProAmpac, Rochester, NY) with 2-6 L of vent fluid for processing shipboard, or to filter between 4.1-6.6 L of fluid through a 142 mm, 0.2 µm PES filter (MilliporeSigma™) for *in situ* samples. Filling and filtering rates ranged between 0.3-1.3 L min^-1^. Fluid was also collected by Niskin bottles mounted on the port forward side of the ROV within the vicinity of the hydrothermal vent, but outside of the range of venting fluid (near vent bottom water) at 2745 m and within the plume by situating the ROV ∼5 m above an active venting site. Background seawater from the water column at ∼2,100 m was also obtained by a Niskin bottle. Upon retrieval, filters from the SUPR sampler were stored in RNAlater™ (Ambion) for 18 hours at 4°C, then moved to -80°C. Niskin samples from the plume and background were emptied into acid-washed cubitainers. Fluids from bags and cubitainers were sampled for prokaryote cell counts by preserving fluid with formaldehyde (1% final concentration). Excess fluid from each bag was also filtered onto sterivex filters (0.2 µm; MilliporeSigma™) and stored with RNAlater identically to the *in situ* filters.

Whenever possible, the same vent fluids and high temperature end-members were also sampled with Isobaric Gas Tight samplers (48) for geochemical analyses, which were processed immediately after recovery of the ROV. Shipboard analyses included pH measured at room temperature (25°C) using a Ag/AgCl combination reference electrode, dissolved H_2_ and CH_4_ by gas chromatography with thermal conductivity detection following headspace extraction, and total aqueous sulfide (∑H_2_S = H_2_S + HS^-^ + S^2-^) following aqueous precipitation as Ag_2_S for subsequent gravimetric determination in a shore-based laboratory. Aliquots of fluid were stored in 30 ml serum vials and acid-cleaned Nalgene bottles for shore-based measurement of total dissolved carbonate (∑CO_2_ = H_2_CO_3_* + HCO_3_^-^ + CO_3_^2-^) by gas chromatography and Mg by ion chromatography, respectively.

### Grazing experimental procedure

Stocks of Fluorescently-labeled Prey (FLP) were prepared using a modified protocol from (49) with monocultures of *Hydrogenovibrio* (Strain MBA27; 50); preparation of FLP prey analog is described in the *Supplementary Text*. The prey type was specifically chosen as a hydrothermal vent representative isolate and was found to have a similar size and morphology to resident bacteria (Figure S1). FLP stained with 5-(4,6-dichlorotriazin-2-yl) aminofluorescein (DTAF) are non-toxic to consumers, and upon ingestion, the DTAF label disappears (49).

Grazing experimental setup and execution followed the guidelines in Caron (51) with modifications described below (Figure S2). A summary of grazing experiments including: site, vent name, depth, incubation temperatures, start times, and sampling time points can be found in Table S1. Vent fluid collected from gas-tight bags was first filtered through 300 µm mesh to remove large multicellular metazoa and transferred into acid-washed and clean 500 ml plastic bottles using a peristaltic pump. Controls were prepared by filtering the fluid through a 0.2 µm filter to ensure that the FLP tracer did not disappear over the course of the experiment in the absence of grazers. Experiments were conducted in duplicate or triplicate and controls were conducted in duplicate (Figure S2). FLP were added at concentrations 50% greater than the *in situ* microbial population, as there were no prior estimates of microbial concentrations or the ability to count cells onboard before the initiation of the experiments Table S1). The suggested amount of labeled prey to be added is between 1 and 10% of in situ microbial concentration (51), thus the higher amount given during these incubations have probably led to overestimation of the estimated rates. Samples at T_0_ were collected for cell counts following addition of FLP and gently mixing by fixing 10 ml of fluid in cold formaldehyde at a final concentration of 1%. Collected fluid in bags (vents) or Niskin bottles (background) remained on the ROV for several hours before the start of each shipboard incubation, thus to minimize additional temperature changes and keep incubation conditions consistent between all experiments, bottles were placed in a dark cooler for incubation, where temperatures ranged between 12-17°C (Table S1). We acknowledge that incubation temperatures may have under (lower temperature compared to *in situ*) or over (higher temperature compared to *in situ*) estimated grazing rates, especially in the experiments using the background sea water where the in situ temperatures were around 1°C.

Grazing incubations were run for a total of 24-48 hours, where sample fluid for FLP counts was preserved with formaldehyde at two time points (T_1_ and T_2_, Table S1). To assess the composition of protistan grazers in grazing incubations via molecular analysis, samples at each time point were vacuum filtered onto 0.2 µm PES filters (Millipore™ Express), stored with RNAlater at 4°C overnight and frozen at -80°C. These were collected in duplicate when possible, and the volume filtered ranged from 0.9-2.7 L (Table S1, Figure S2). In some cases, a T_0_ sample was taken after addition of FLP and before incubation started, providing an assessment of the degree to which the community composition in the initial water samples was altered by sample handling between collection in situ and initiation of the experiments shipboard. A molecular sample at T_0_ was not always collected (*e*.*g*., Candelabra and Sir Ventsalot), and a subset of grazing experiments were conducted at different time points ranging from 18 to 36 hours (Table S1).

To track the disappearance of FLP over time, triplicate slides were prepared from each time point in a shore-based laboratory by filtering 2-4 ml of preserved fluid from each time onto 0.2 µm black polycarbonate filters. Following filtration, 10-15 µl of a stain solution made with 4′,6-diamidino-2-phenylindole (DAPI; ∼10 µg/ml; see *Supplementary Text*) was gently pipetted onto the filter, and covered with a cover slip. Experimental and control slides were counted using epifluorescence microscopy within 1-2 days and stored at 4°C. FLPs were counted under the fluorescein isothiocyanate (FITC) filter at 100x or 63x; 16 fields of view were counted and the cell ml^-1^ concentration was determined from this value for each slide. The technical error rate was calculated by taking the percentage of the standard deviation over the mean for replicate counts. This technical error rate percentage was used to set the threshold at which a change in FLP abundance over time was considered true (*e*.*g*., if the percent change in FLP from T_0_ to T_1_ did not exceed the technical error rate, the loss of FLP by T_1_ was not considered significantly different from T_0_).

The concentration of FLPs at each time point was averaged across replicates. The difference in FLP concentration (cells ml^-1^) from T_0_ and T_F_ was used to estimate the number of cells grazed (G). For each experiment, T_1_ or T_2_ was chosen as T_F_, when the loss in FLP exceeded the technical error rate. In the case where both T_1_ and T_2_ exceeded the range of error, T_1_ was chosen as T_F_. Using a model described in Salat and Marrasé (52) the number of cells grazed by protists (*G*) was estimated using the equation:

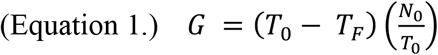

where T_0_ and T_F_ equal the average FLP concentrations at the beginning and end of the experiment and N_0_ equals the concentration of *in situ* prokaryote cell concentration (52). This model assumes that the ratio of FLP to *in situ* microbial prey remains consistent. The grazing rate was calculated by normalizing *G* to time at T_F_ (cells ml^-1^ hr^-1^). The daily prokaryote turnover percentage was calculated by multiplying the *in situ* prokaryote cell concentration (taken at T_0_) by the estimated grazing rate per day (9, 52). Grazing rates were converted to measures of carbon biomass using the assumption that the amount of carbon per prey cell is 86 fg C (Morono *et al*. (27); Table S1).

### Eukaryotic molecular sample processing

Samples collected for molecular analyses included *in situ* filters from the SUPR sampler, shipboard sterivex filters, or time points from grazing experiments (all collected into 0.2 µm pore size filters; MilliporeSigma™). RNA was extracted and the 18S rRNA gene was reverse transcribed; the V4 hypervariable region (53) was amplified according to Hu *et al*. (54); protocols.io; dx.doi.org/10.17504/protocols.io.hk3b4yn) and as described in the *Supplementary Text*. Samples were multiplexed, pooled at equimolar concentrations and sequenced with the MiSeq 300 x 300 bp PE kit at the Marine Biological Laboratory Bay Paul Center sequencing facility.

### Prokaryotic sample processing

Prokaryotic cells were enumerated in formaldehyde-fixed fluids using DAPI (see *Supplementary Text*). DNA was extracted from PES filters or sterivex filters (0.2 µm) as described in (55) and the *Supplementary Text*. 16S rRNA gene amplicon libraries were prepared and sequenced by the UConn Microbial Analysis, Resources, and Services using modified EMP 16S rRNA gene V4 primers 515F and 806R (56–59).

### Sequence analysis

All sequences were quality controlled and processed in QIIME2 (v2019.4; 60). Chimeric sequences were removed (pooled) and Amplicon Sequence Variants (ASVs) (61). ASVs from 18S rRNA amplicons were assigned taxonomy using the Protist Ribosomal 2 Database (v4.12; https://github.com/pr2database/pr2database;(62). Taxonomy assignment was performed with the naive Bayesian classifier method in the DADA2 R package with a minimum bootstrap of 70 (61, 63). Removal of contaminant 18S rRNA sequences is described in the *Supplementary Text*. For 16S rRNA gene derived ASVs, the SILVA database (v132; 64) was used for taxonomy assignment.

Molecular samples from *in situ* filters and shipboard sterivex filters were treated as replicates, where sequence counts were averaged across replicates at the ASV level. ASV taxonomy assignment for both the 18S rRNA and 16S rRNA gene was manually curated and visualized to highlight the main taxonomic groups (see *Supplementary Text*). Due to the compositional nature of tag-sequence datasets, data were transformed by center-log ratio ahead of Principle Coordinate Analysis and to visualize ASV-level changes across samples (65, 66).

To detect possible microbial interactions, Sparse InversE Covariance estimation for Ecological Association and Statistical Inference (SPIEC-EASI) analysis was performed using the cross-domain approach with ASVs from 18S rRNA and 16S rRNA gene results (67). SPIEC-EASI is designed to minimize spurious ASV-ASV interactions that result from the influence of compositional nature of tag-sequencing results (46). Only *in situ* samples that were found in both the 18S rRNA and 16S rRNA gene amplicon results were considered for the network analysis. Both datasets were subsampled to include ASVs that appeared in at least 3 samples, had at least 50 sequences each, and made up at least 0.001% of the sequenced reads. 18S rRNA and 16S rRNA gene datasets were each center-log ratio transformed then SPIEC-EASI was run using the Meinshausen-Buhlmann’s neighborhood selection estimation method. Significant interactions to infer putative predator-prey relationships were determined by subsetting only interactions between 18S rRNA and 16S rRNA-derived ASVs.

### Data availability

A complete compilation of code to reproduce all analyses is available at https://shu251.github.io/protist-gordaridge-2021/. A GitHub repository also includes raw microscopy count results, raw sequence count information, and ASV tables. Both 18S rRNA and 16S rRNA amplicon sequences have been deposited in the Sequence Read Archive under BioProject PRJNA637089 (Table S2).

## Acknowledgements and funding sources

Authors would like to thank Chip Breier and Darlene Lim for their contributions to the field operations. We would also like to extend gratitude to David Caron, Paige Hu, Susanne Menden-Deuer, Roxanne Beinart, and Alexis Pasulka for helpful conversation and discussion regarding grazing experiments and deep-sea protistan diversity and to Jesse McNichol for discussion of carbon fixation rates. Gretta Serres and Patrick Carter provided laboratory support for preparation of FLP. This research was supported by the NASA Planetary Science and Technology Through Analog Research (PSTAR) Program (NNH16ZDA001N-PSTAR) grant (16-PSTAR16_2-0011), NOAA Office of Ocean Exploration and Research, Ocean Exploration Trust and NOAA-OER grant NA17OAR0110336. This research used samples and data provided by the Ocean Exploration Trust’s Nautilus Exploration Program, Cruise NA108. EH was supported by the WHOI Summer Student Fellowship and NSF REU (OCE-1852460). The NSF Center for Dark Energy Biosphere Investigations (C-DEBI) supported JAH as well as SKH through a C-DEBI Postdoctoral Fellowship (OCE-0939564). This is SUBSEA Publication Number SUBSEA-2021-xxx and C-DEBI contribution number XXX.

## Figure Legends

**Figure S1**. (**A-C**) Location of Sea Cliff and Apollo vent fields on the Gorda Ridge shown by the pink and yellow stars in **C**, respectively. Map modified from Clague *et al*. (21). (**D**) Images taken during sample collection from the four vent sites where grazing experiments were conducted. Copyright Ocean Exploration Trust, Inc. (**E**) Epifluorescence image from grazing experiment counts where solid arrows denote *in situ* bacteria and dashed arrows indicate fluorescently-labelled prey (FLP). The image demonstrates that the FLP were within the size range of the *in situ* microbial population. (**F-G**) Images taken of eukaryotic cells from the grazing experiments. Preserved sample material was set aside and filtered onto 0.8 µm filters to image vent associated protists under epifluorescence.

**Figure S2**. Schematic showing sample origin and general processing steps for molecular survey and grazing experiments. ROV Hercules collected *in situ* filters and fluid from diffuse fluid, plume (a couple of meters above the venting fluid), near vent bottom water, and background seawater.

**Figure S3**. Loss in fluorescently-labeled prey (FLP) over time for each grazing experiment. FLP loss is shown for both the controls (left panel) and experimental treatments (right panel). Error bars represent the standard mean error and data points represent the average of replicate samples. Shaded area represents the determined microscopy error percentage above and below the T_0_ time point; this metric serves to demonstrate significant changes in FLP concentration. FLP change in the control experiments (right) demonstrates that introduced FLPs did not disappear for other reasons besides grazing; for most experiments, FLP concentration in the control samples remained within the margin of error at the determined T_F_ time point (left). Experiments where FLP loss at T_1_ or T_2_ fell below the microscopy error were used to calculate the extent and rate of grazing; labeled with an asterisk (*). See Materials and Methods for additional explanation.

**Figure S4**. Relationship between measurements of protistan grazing pressure (top to bottom) and environmental parameters (from left to right): temperature, prokaryote concentration percent seawater of diffuse fluid, pH, and magnesium. Environmental parameters (x-axis) are shown in relation to (**a**) grazing rate in cells ml^-1^ hour^-1^, (**b**) prokaryote turnover percentage day^-1^, and (**c**) µg C L^-1^ day^-1^ consumed. Dashed lines represent slope from linear regressions and r^2^ values are listed in the top left corner of each plot. Environmental data were obtained from SUPR bag samples that were used for grazing experiments (Table 1). Estimates of grazing pressure (grazing rate, rate of prokaryote turnover, and rate of carbon consumed) do not strongly correlate with environmental parameters, with the exception of prokaryote turnover percent day^-1^ with temperature.

**Figure S5**. (**a**) Taxonomic breakdown of samples from the Gorda Ridge collected from background seawater, diffuse hydrothermal fluids, and associated grazing incubation experiments. Bar plot shows the relative sequence abundance for each sample (including replicates) and colors designate major protistan taxonomic groups, which has been manually curated (see *Materials and Methods*). Bar plot also varies from the main text by showing the relative sequence abundance of opisthokonta and unassigned sequences. (**b**) Average hierarchical clustering of all samples following relative abundance transformation and dissimilarity calculation. Colors and symbols are consistent with Figure 2.

**Figure S6**. (**a**) Taxonomic breakdown of 16S rRNA gene amplicon results from Sea Cliff and Apollo hydrothermal vent fields derived from background and *in situ* vent samples. Colors denote bacteria or archaea taxonomic groups. Most samples are the result of averaging among 2 or 3 samples taken at the same time. The two leftmost samples originated from background shallow (150 m) and deep seawater (>2000 m). The “Other” category represents 16S rRNA gene-derived ASVs that were less than 0.1% in abundance or were manually removed due to known extraction kit representatives. (**b**) Ordination analysis of all samples, including replicates, from the 16S rRNA gene-derived sequence data. Data was center log-ratio transformed ahead of PCA analysis, similar to Figure 2b for the 18S rRNA-based ordination analysis.

**Figure S7**. Distribution of protistan ASVs among background and vent sites (see main text for description of cosmopolitan versus resident). Here, the resident and cosmopolitan populations are further grouped by the presence of ASVs throughout the Gorda Ridge. Relative abundance of (**a**) sequences and (**b**) ASVs based on the distribution of ASV occurrence (denoted by color). Total number of (**c**) sequences and (**d**) ASVs based on the distribution of ASV occurrence.

**Table S1**. Complete experiment details for each grazing incubation. Sample information (orange) rows list dive IDs and identifiers designated from EV Nautilus. Grazing incubation details (green) list start times and sampling time points (two per experiment) for each experiment and temperature of incubations. Due to the novel nature of these experiments, incubations were run at a variety of times, but were sampled at either approximately T_18_ and T_24_ or T_24_ and T_36_. Finally, the bottom columns (blue) list which time point was found to have a significant difference from the T_0_ (based on microscopy error; see *Materials and Methods* and Figure S3), the fluorescently-labeled prey (FLP) cell concentration at T_0_ and T_F_, the average *in situ* prokaryote cell concentration, calculated mortality factor (m), grazing rate, and estimated prokaryote turnover percentage, results from carbon conversion estimates, and associated statistics.

**Table S2**. Sample names, metadata, and SRA IDs for all sequence samples in this study. Both 18S and 16S rRNA amplicon sequencing was conducted for this study, all sequences are submitted under SRA BioProject PRJNA637089.

**Table S3**. Total number of ASVs (top table) and sequences (bottom table) for each major protistan group. Columns list each sample type and represent the average across replicates and the sum across samples from the same grazing incubation.

**Table S4**. Total number of ASVs (top table) and sequences (bottom table) for each protist group at the class or family level. Taxonomic levels were curated to the class or family level as shown in Figure 3. Columns list each sample type and represent the average across replicates and the sum across samples from the same grazing incubation.

**Table S5**. The 10 most abundant ASVs within each protistan taxonomic group. Feature.ID reports the unique ASV identification, Distribution indicates if the ASV was found to belong to the resident or cosmopolitan population, and other columns report the taxonomic classification. The ASV size is the total number of sequences associated with the ASV.

**Table S6**. Summary of the significant 18S-16S ASV correlations derived from SPIEC-EASI results. The first four columns report the ASV Feature.ID and complete taxonomic name for the 18S and 16S ASV involved in the putative interaction. The following columns report the 18S ASV distribution and broader taxonomic classifications for both the 18S and 16S results. Finally, the weight reports the correlation value between the inferred relationships, the value reflects the strength of the interaction.

