## Supplementary Information for "Protistan grazing impacts microbial communities and carbon cycling at deep-sea hydrothermal vents"

### Supplementary Materials and Methods

*Preparation of analog prey*

Stocks of Fluorescently-labeled Prey (FLP) were prepared using the protocol from (1) with some modifications. Several monocultures of *Hydrogenovibrio* (Strain MBA27; (2)) were grown in Luria Broth Base media (LB broth; 5 g tryptone, 2.5 g yeast extract, 5 g NaCl in 500 ml of Milliq water, at pH 7.0 and autoclaved). To avoid clumping or biofilm formation, 70 ml of culture was grown in 250 ml flasks at 35ºC on a shaker plate (115 RPM) overnight. To heat-kill and stain the bacterial prey, 35 ml volumes of culture were filtered through a 20 µm mesh into 50 ml falcon tubes. These falcon tubes were centrifuged at 7,500 RPM for 30 minutes to form a pellet of cells. The liquid was decanted from each tube and the pellet was resuspended in 20 ml of Sea Salt Broth (SSB) by vortexing vigorously for 10 minutes. SSB (5X) was prepared by mixing and autoclaving: 98g NaCl, 16.5g $Na_2SO_4$, 1.5g KCl, 0.25g KBr, 0.1g $H_3BO_3$, 0.1g and 44g of $MgCl_2$-$6H_2O$ into 1 liter of sterile Milliq water. SSB rinse steps were repeated three times to remove away additional LB broth, by repeatedly resuspending cells in 10 mL of SSB. 150-µl of DTAF (filter sterilized stock of 5-(4,6- dichlorotriazin-2-yl) aminofluorescein) was added to each resuspension and vortexed vigorously for 10 minutes. All tubes were then incubated in a water bath (60ºC) for 2 hours; every 10 minutes during this incubation each tube was vortexed vigorously for 30 seconds to reduce cell clumping. Following incubation, 25 ml of SSB was added to each tube and centrifuged at 7,500 RPM at 15ºC for 20 minutes to pellet the heat-killed stained cells. Excess liquid was removed and replaced with 10 ml of SSB; tubes were vortexed for 10 minutes and an additional 25 ml of SSB was added before spinning again at 7,500 RPM for 20 minutes. These wash steps were repeated 3 times to rinse away excess stain. Cells were concentrated by resuspending in only 10 ml of SSB after the final wash step. Resuspended, stained cells were combined into a single flask and continually vortexed while aliquoting 1 ml volumes into cryovials. Cryovials containing FLP stock were frozen and stored at -80ºC. FLP stock concentration was estimated by preparing a slide from randomly chosen cryovials and counting under epifluorescence microscopy.

*Cell enumeration*

Fluid from each study site or grazing experiment was preserved with formaldehyde (1% final concentration) for downstream cell enumeration. For each grazing experiment, 2-4 ml of fluid was filtered and 1-2 ml of fluid or background seawater was filtered to count prokaryotic cell concentration. Preserved fluid was filtered onto 0.2-µm black polycarbonate filters (25 mm size; PCTE, Sterlitech PCTB0225100) with a GF/F backing filter (25 mm; Whatman) with a peristaltic pump. Following filtration, filters were transferred to glass slides and let to dry in the dark. Then a stain solution was added to each filter and covered with a plastic coverslip; the stain solution consisted of 137.5 µl of Citifluor, 50 µl TE buffer, 50 µl of Vectashield, 25 µl of PBS (1X), and 2.5 µl of 4′,6-diamidino-2-phenylindole (DAPI; 1 mg/ml concentration; Sigma D9542). The stain solution stained prokaryotic cells with DAPI, which were subsequently imaged with epifluorescence microscopy (Axio 2 Imager, Zeiss) under blue-light excitation (365 nm). To enumerate FLP in the grazing experiments, DTAF-stained cells from FLP excited under FITC filter and were thus differentiated from the *in situ* prokaryotic cells. Slides dedicated for *in situ* prokaryotic cell counts were counted under 100X with a minimum of 15 fields of view.

*Extraction of eukaryotic genetic material*

Frozen filters were thawed and placed into sterile 15 ml falcon tubes with sterile forceps, and 1-2 mL of RNeasy Lysis Buffer (RLT with β-Mercaptoethanol, Qiagen, Valencia, CA, USA) and RNase-free silica beads were added to each tube. Falcon tubes were bead-beaten by vortexing vigorously for 5 minutes. The original sample collection tubes with RNAlater were centrifuged to pellet any cellular material left in the RNAlater; the RNAlater was removed and replaced with 500-ul of RLT buffer. This was vortexed and added to the 15-ml falcon tube. RNA was extracted with the Qiagen RNAeasy kit (Qiagen #74104) with the in-line genomic DNA removal step (RNase-free DNase reagents, Qiagen #79254). RNA concentrations were determined using the Quant-iT RiboGreen RNA assay kit (ThermoFisher Scientific). Extracted RNA was reverse transcribed into cDNA using a cDNA synthesis kit (iScript Select cDNA Synthesis, BioRad, #1708896, Hercules, CA); the concentration of RNA was normalized for the cDNA synthesis reaction (0.4 ng of RNA). Primers targeting the V4 hypervariable region of the 18S rRNA gene (3, 4) were used in PCR reactions, which consisted of a final concentration of 1X Q5 High Fidelity Master Mix (NEB #M0492S, Ipswich, MA), 0.5 µM each of forward and reverse primers, and 1 ng of genetic material. The PCR thermal protocol started with an initial activation step (Q5 specific) of 98°C for 2 min, followed with 10 cycles of 98°C for 10 s, 53°C for 30 s, 72°C for 30 s, and 15 cycles of 98°C for 10 s, 48°C for 30 s, and 72°C for 30 s, and a final extension of 72°C for 2 min (modified from Rodriquez Martinez et al. 2012). The original extract total RNA was also PCR amplified to ensure no genomic DNA was present in the sample. PCR products were checked by confirming the presence of an ~400 bp product on an agarose gel. In cases with no amplification, the PCR reaction was repeated with a higher concentration of cDNA (1.5-2 ng). If this did not yield the expected PCR product, the reaction was repeated with an additional 5 cycles. Three shipboard blanks (MilliQ water) and one extraction blank were also extracted and PCR amplified; while no PCR product was observed in these control samples they were processed identically to all samples and sequenced. All PCR products were cleaned using the AMPure bead clean up (Beckman Coulter #A63881, Brea, CA).

*Extraction of prokaryotic genetic material*

DNA was extracted from filtered vent fluids on 142 mm 0.2 µm filters (PES MilliporeSigma™) or sterivex filters (0.2 µm pore size), which had been stored in RNALater. Sterile forceps and ethanol flamed scissors were used to unravel and cut the filter (1/3rd of the filter was used for each extraction). For sterivex filters, the entire sterivex cartridge was opened. Filters were rinsed twice in sterile PBS and cells were pelleted by centrifugation. DNA extraction buffer (50 mL, 100mM Tris, 100mM EDTA, 100mM NaH2PO4, 1.5M NaCl, 1% CTAB), 20 µl proteinase K (10mg/mL), and 40 µl of lysozyme (50 mg/mL) was added to tubes with pelleted cells and filters and put through three  free-thaw cycles. Filter sterile SDS was added to each tube and incubated for 2 hours at 65C. DNA extract was isolated by isolating the organic phase following a phenol:chloroform:isoamyl alcohol (25:24:1, pH 8.0) extraction (x2) and DNA was precipitated by mixing with 100% isopropanol and incubating overnight at room temperature, then DNA was isolated and washing with ice cold 70% ethanol (x3). DNA extracts were quantified with the Quant-iT PicoGreen dsDNA assay kit (ThermoFisher Scientific).

*Sequence analysis*

All sequences were processed through a snakemake pipeline available on Github:

https://github.com/shu251/tagseq-qiime2-snakemake. This pipeline processes raw sequences through fastq and multiqc, trims low quality and adapter reads, and executes all steps in the QIIME2 pipeline.

Most samples are the result of averaging among 2 or 3 samples taken at the same time (see Figure S5). The two leftmost samples originated from background shallow (150 m) and deep seawater (>2000 m). Data not shown include ASVs identified as opisthokonts or left unassigned (see Figure S5 and Tables S3 and S4). The choice to exclude opisthokonts from the majority of analyses is three fold: prefiltering of samples for incubations would have removed a subset of multicellular metazoa, sequence database and processing is most suitable for protists, and opisthokonts were not the focus of this study. The 'Unassigned-Eukaryote' category represents ASVs left without a taxonomic assignment. 'Other' classifications for each group represent ASVs either made up less than 0.1% of the groups sequences, or were only assigned to the Supergroup or Phylum level.

To remove contaminant sequences (based on shipboard and lab blank samples) the R packages 'decontam' and 'phyloseq' were used (5, 6) in R v3.6.1 (7). A threshold of 0.5 was used to compare the prevalence of ASVs in control samples versus environmental samples; ASVs that were more prevalent in the control samples compared to the environmental samples were considered contaminants and removed from the dataset; control samples included shipboard MilliQ water filtered at the time of sampling and extraction blanks prepared during the nucleic acid extraction. This approach was conducted for only 18S rRNA gene results.

### Supplementary Results

*Geochemistry at Sea Cliff and Apollo*

Low-temperature diffusely venting fluids were collected at the Sea Cliff and Apollo hydrothermal vent field along the Gorda Ridge (Figure S1; 8). The concentration of bacteria and archaea was 0.5 - 1 x $10^5$ cells $ml^{-1}$ in low temperature vent fluids, compared to background seawater concentrations of 3 - 5 x $10^4$ cells $ml^{-1}$ (Table 1). The low-temperature Candelabra and Sir Ventsalot sites were situated close to the sites of high temperature venting fluid and had similar geochemistry to one another. These high temperature vents (Candelabra, 298ºC and Sir Ventsalot, 292ºC) were also sampled to determine end-member geochemistry of the venting fluids, as part of our larger SUBSEA study (9). All diffuse vents sampled in both fields represented a mixture of this high temperature vent fluid with seawater (10). At both locations, high temperature fluids were acidic (pH 2.8-4.5 measured at 25ºC), with near-zero magnesium concentrations (2.1-2.5 mM) and hydrogen sulfide concentrations typical of mid-ocean ridges, 2.5-3 mM (Table 1). Hydrogen concentrations ranged from 62-71 µM and methane concentrations ranged between 66-68 µM. During sample vent fluid sample collection (30-40 minutes) the fluids being sampled fluctuated between 3-72ºC, due to mixing (Table 1). Mixed fluids at Candelabra and Sir Ventsalot were determined to contain 88% and 98% seawater, respectively (Table 1). The temperature maxima at Mt. Edwards and Venti Latte were lower and ranged from 11-40ºC; these sites also had visible tube worm clusters (*Paralvinella palmiformis*; Figure S1; Table 1). While Venti Latte was 97% seawater, Mt. Edwards was 82% (Table 1). A maximum hydrogen concentration of 127 µM was detected at Mt. Edwards vent, whereas hydrogen was undetectable at Venti Latte, and methane concentrations ranged from 0.9-10 µM in the diffuse vent fluid (Table 1).

*Grazing experiment estimations*

Despite technological challenges in the present experiments, trends in grazing rates and variation among estimated grazing rates were independent of experimental design details. Each experiment conducted at Gorda Ridge demonstrated measurable loss in the introduced Fluorescently-labeled Prey (FLP; Figure 1a; Figure S3).

The number of bacteria grazed (*G*) during the incubations was estimated using Models I and II from (11). Based on Model I, *G* was ~8,900 in the near vent bottom environment and ranged between ~16,800 - 32,900 at the vent sites (see Table S1). Model II values were slightly higher, but comparable to the results from Model I (<30% different; Table S1). Model II incorporates the initial and final concentration of natural bacteria, yet the natural bacteria population at $T_F$ was not collected. Therefore, as assumption of our experiments is that the proportion of analog prey (FLP) with respect to the natural bacterial population did not change during the incubation period (meaning, growth of natural bacteria is negligible). Model I was chosen for the main analysis in this study, as this is an assumption of Model I. Model III was not incorporated as the input values for FLP and natural bacteria population would not have differed from Model II.

Prokaryotic turnover percentage $day^{-1}$ at each of the vent sites was 32.7% at Mt. Edwards, 28.1% at Venti Latte, 42.6% at Candelabra, and 62.1% at Sir Ventsalot (Figure 1c; Table S1). Using a carbon conversion factor of 86 fg carbon $cell^{-1}$ (12), estimated carbon consumption rates were 0.53 µg of carbon $L^{-1}$ $day^{-1}$ in the near vent bottom seawater and 1.45 µg of carbon $L^{-1}$ $day^{-1}$ at Mt. Edwards, 1.86 µg of carbon $L^{-1}$ $day^{-1}$ at Candelabra, 3.40 µg of carbon $L^{-1}$ $day^{-1}$ at Venti Latte, and 3.77 µg of carbon $L^{-1}$ $day^{-1}$ in diffuse vent fluids (Table S1).

Grazing factor (*G*) determined using Model I in cells $ml^{-1}$ consumed $day^{-1}$ was multiplied by a carbon conversion factor to estimate µg C $L^{-1}$ $day^{-1}$. A carbon conversion factor of 86 fg C $cell^{-1}$ from (12) was used in the main text. Estimated carbon consumed using a carbon conversion value of 173 fg C $cell^{-1}$ is also reported in Table S1 (13, 14). Calculations for all grazing estimates and estimates of consumed carbon are available at: https://shu251.github.io/protist-gordaridge-2021/.

*Amplicon sequencing*

A total of 1.43 million 18S rRNA amplicons and 1.08 million 16S rRNA gene amplicons were sequenced and designated as ASVs (passing sequence quality control; see Methods). There were a total of 9028 and 6497 ASVs determined from the microbial eukaryotic and prokaryotic results, respectively. Additional quality control accounted for the composition and relative abundance of ASVs found in shipboard blanks and extraction control samples for the 18S rRNA gene results. 34 ASVs were found to be likely contaminants and removed; this removed only 1.24% of the sample reads (Figure S3). A background sample from the 18S rRNA results was removed due to low quality sequencing (BSW020). After averaging across replicates, there was a total of 9028 ASVs and 1.43 million reads in the eukaryotic dataset. Samples from the same vent site that originated from separate filters collected were considered biological replicates. Results from biological replicates were found to mainly group together (Figure S5); thus the majority of downstream visualizations show data averaged across replicate samples, when replicates are shown or were considered during analysis it is noted. ASVs assigned to opisthokonts, which comprised 12.9% of all 18S sequence data (615 ASVs), were not the direct focus of this study; thus the majority of these ASVs were not considered in downstream analyses. A summary of opisthokont diversity is reported in Figure S5. ASVs where the taxonomic assignment was "Eukaryote" were considered "Unassigned"; there were 1058 unassigned ASVs, which made up 2.8% of all 18S sequence data. For cluster dendrograms and ordination analysis, opisthokont and unassigned reads were not removed. Before further analysis, 16S rRNA gene results identified as eukaryotic or Unassigned were removed. This left a total of 1.08 million sequences and 6497 ASVs from the prokaryotic tag-sequencing results that were used in downstream analyses.

*Detailed characterization of microbial diversity*

The diversity and distribution of the vent-associate protistan community was determined from 18S rRNA gene tag-sequencing.The majority of protists detected in both *in situ* and grazing experiment samples and found to be associated with 16S rRNA gene ASVs were primarily made up of known heterotrophic species. All vent sites were characterized by high relative abundances of ciliates, dinoflagellates, and Syndiniales; together these three alveolate groups represented 50% or more of the 18S rRNA reads in most samples (Figures 2a, S5; Table S3). Ciliates had high relative abundances at all vent sites compared to plume and background samples, which had higher relative abundances of stramenopiles. After alveolates, rhizaria and stramenopiles were the next two most numerous groups detected at the vent sites. Radiolaria were also detected in all samples, with the highest relative abundance within the Mt. Edwards and Candelabra plume samples (Figure 2a). The background, plume, and near vent bottom water bacteria and archaea community composition based on 16S rRNA genes was also distinct from the community detected at each vent site (Figure S6a).

Ordination results demonstrated that the microbial eukaryotic community clustered primarily by location, and then by sample type (compare color vs. symbol in Figure 2b), with some exceptions. The shallow water sample clustered separately from all other samples, while the deep seawater background samples clustered with Sir Ventsalot and Mt. Edwards. Replicate samples from both 18S rRNA and 16S rRNA gene results were similar to one another (identical symbol and colors in Figures 2b, S6). For archaea and bacteria, samples from Venti Latte, Mt. Edwards vent, and near vent bottom water clustered closely to one another, whereas Sir Ventsalot and Candelabra vents showed more variability (Figure S6). Background deep seawater samples clustered with the vent plume and near vent bottom water samples (Figure S6).

A network analysis from the 18S and 16S rRNA gene tag-sequence results was conducted to determine if significant interactions may be indicative of predator-prey relationships. This was conducted with a subset of 207 eukaryotic and 158 prokaryotic using SPase InversE Covariance Estimation for Ecological Association Inference (SPIEC-EASI); SPIEC-EASI is a computational tool that constructs a network based on co-occurring ASVs to infer an ecological association, while minimizing the negative impacts of compositional sequence datasets (15). 537 protist-bacteria and protist-archaea significant interactions were recovered (Table S6). There was a higher total number of significant interactions between protists and prokaryotes among the cosmopolitan protist populations, where interactions involving dinoflagellate and ciliate predators and prokaryotic ASVs affiliating with *Thaumarchaeota-Nitrososphaeria*, *Proteobacteria-Gammaproteobacteria* occurred most frequently (Table S6; Figure 4a). Among the resident protistan population, interactions between ciliates and *Proteobacteria- Alphaproteobacteria, Gammaproteobacteria*, *Thaumarchaeota-Nitrososphaeria*, or *Epsilonbacteraeota-Sulfurimonas* were the most abundant interaction (Figure 4b). In total, protistan groups with the highest number of significant interactions with potential prey populations included ciliates (133 ASVs), dinoflagellates (112 ASVs), Syndiniales (82 ASVs), radiolaria (68 ASVs), and MArine STramenopile (MAST) groups (36 ASVs) (Table S6; Figure 4). 16S rRNA gene-derived ASVs with the highest number of significant interactions with eukaryotic taxa included the *Proteobacteria-Alphaproteobacteria* (89 ASVs), *Thaumarchaeota-Nitrososphaeria* (78 ASVs), *Proteobacteria-Gammaproteobacteria* (74 ASVs), and *Epsilonbacteraeota-Sulfurimonas* (54 ASVs) (Table S6; Figure 4).

Within each major protistan taxonomic group, "Other" categories or taxonomic groupings that did not resolve beyond the phylum level represent sequences without a close representative in current databases. Many of these ASVs were associated with the resident vent population and therefore represent yet to be recovered diversity at deep-sea hydrothermal vents (Tables S3-S5).

*Ciliates*

The consistently higher relative abundance of ciliates at the vent sites and grazing incubations, relative to background and plume, revealed the group to be the predominant protistan grazer. Among the highly abundant ciliate groups, Spirotrichea-Strombidiida, Spirotrichea-Choreotrichida, Heterotrichea, and Oligohymenophora ciliates were detected in both cosmopolitan and resident populations. The most abundant Spirotrichea-Strombidiidia ASVs included members of the *Tontoniidae* family, including *Spirotontonia*, *Pseudotontonia simplicidens, Laboea strobila,* and *Varistrombidium kielum* (Table S5). The Strombidiida order was also found to have the highest total number of significant interactions with 16S ASVs (n=47), together with their broad distribution, this suggests that many species may be non-specific opportunistic grazers. *Strobilidiidae* and *Leegaardiellidae* were the most abundant members of the Choreotrichida detected, the latter of which has been reported from at East Pacific Rise and within vent-influenced plumes in the Okinawa Trough (16, 17).

Members of the ciliate Heterotrichea class may have outcompeted other protists with the grazing incubations, as evidenced by the relative increase in sequences in the Mt. Edwards and Sir Ventsalot grazing experiments compared to *in situ* and $T_0$. Heterotrichea ciliates were dominated by ASVs belonging to *Folliculinidae* in both ubiquitous and resident populations (Table S5). *Folliculinidae* ciliates are known vent endemic species that form sessile colonies on hard substrates (18, 19) and have been found at vents throughout the NE Pacific (20, 21) and at methane seeps (22). Blue-purple mats of Folliculinid ciliates were observed during ROV operations (Figure S1b), confirming their presence at Gorda Ridge; yet, their abundance in the *in situ* and grazing experiment samples is likely derived from their motile free-living stage that is non-feeding (18). Stable isotope experiments with Folliculinid ciliates have shown that food sources are spatially variable and correspond to their proximity to venting fluid (20, 22). Folliculinid ciliates primarily depend on symbiotic sulfide-oxidizing bacteria and in this study were not found to significantly interact with any 16S ASVs. This demonstrates the value in our approach to compile paired 18S and 16S sequence datasets with quantitative measurements of grazing to investigate food web dynamics.

In addition to Spirotrichea-Strombiidiida, resident ciliate classes were largely comprised of Spirotrichea-Euplotia, Spirotrichea-Choreotrichida, Oligohymenophorea, and Litostomatea (Figure 3, Table S4). Within the Euplotia class the most abundant ASVs belonged to the *Aspidiscidae* or *Uronychiidae* families, including *Aspidisca*, *Uronychia setigera*, and *Paradiophrys irmgard* (Table S5). The Oligohymenophorea group ASVs were dominated by scuticociliates, notably the *Philasterida* family, and the Litostomatea were primarily composed of species belonging to *Pleurostomatida* or *Lacrymariida**e*** (Table S5). *In situ* ciliate diversity among the resident population varied with respect to hydrothermal vent site, which was similar to the relative abundances of ciliates among the grazing incubations (Figures 3, S5).

*Dinoflagellates & Syndiniales*

Within the dinoflagellates and Syndiniales, most classes were represented in both cosmopolitan and resident populations (Figure 3); exceptions among the dinoflagellates included Torodiniales within the cosmopolitan population, and Suessiales, Gonyaulacales, and Apicomplexa which were classified as resident. Within the Syndiniales, Dino-Groups -I and -II had the highest number of ASVs within the resident population (Tables S4-S5).

*Rhizaria*

Within the rhizaria, the acantharia groups, RAD-B, and RAD-C groups dominated within the plume and near vent bottom water samples, demonstrating that these environmental clades are found throughout the deep-sea, and may be specifically suited to thrive in the seawater-diluted plume environment (Figures 2, 3). The difference between the background and vent-associated cercozoan distribution was specifically distinct, where the vent-only cercozoa were typically identified as Filosa (with the expectation of the Imbricatea order) and the cosmopolitan cerocozoa were found to be Endomyxa (Tables S4-S5).

*Stramenopiles*

The MArine STramenopile (MAST group) ASVs were found in both the cosmopolitan and resident populations (Figure 3). MAST-3 ASVs represented the highest proportion of MAST ASVs (Table S5). Stramenopiles belonging to the flagellated Chrysophyceae group were found in both resident and cosmopolitan populations; resident Chrysophyceae were more abundant and predominantly comprised of *Paraphysomonas foraminifera,* while the majority of cosmopolitan ASVs were identified as environmental clade H (Tables S4-S5). The near vent bottom water samples were overwhelmed by sequences identified as *Caecitellaceae* and *Cafeteriaceae* (Figure 3).

Many species within the stramenopiles (e.g., pelagophytes, bacillariophyta) and hacrobia are more commonly associated with euphotic ecosystems and play important roles as phytoplankton in the upper water column. ASVs belonging to these groups did not make up a large portion of the sequenced data and may have originated from sinking material (23, 24). Further, taxonomic assignment of these ASVs was not resolved at the genus or species level, indicating that sequences may belong to species with no close relatives in the reference database (i.e., deeply branching or novel species).

*Amoebozoa, Excavata, and Apusozoa*

Amoebozoa, excavata, and apusozoa protistan classes were detected only among the resident vent population (except for sequences classified as *Hilomonadea*; Figure 3). Amoebozoa included Lobosa (*Flabellulidae*, *Vannella*) and Breviata (NAMAKO-1), the excavata were further identified as Discoba (Jakobida) or Metamonada (*Carpediemonas*), and apusozoa included Hilomonadea (e.g., *Ancryomonas microns*) and Apusomonadidae Group-I (Table S5). Species belonging to these groups also inhabit other extreme environments, such as microbial mats found in caves and deep-sea abyssal plains or sediments (25–27). However, surveys of their distribution may be limited by their underrepresentation in sequence repositories (28), debated phylogenetic placement, or their overall low biomass keep them below the limit of detection for sequence surveys (29). Of the amoeboflagellates found at Gorda Ridge, abundant ASVs within the Breviata order were similar to putative anaerobic species found in the anoxic sediment of a meriomictic lake (NAMAKO-1; 27). This molecular survey also revealed a excavata ASVs included species identified as *Jakoba libera*, which were found at the site of the lower temperature diffuse venting fluid (Mt. Edwards and Venti Latte). These heterotrophic flagellates were also identified as vent endemics in Mariana Arc, demonstrating that excavata species may represent important vent endemic bacterivores.

*Hacrobia*

The majority of the hacrobia sequences and ASVs were classified as cosmopolitan and identified as haptophytes or cryptophytes (Tables S3-S4, Figure 3). While vent resident hacrobia ASVs did not make up a majority of the sequenced reads, resident hacrobia classes included Telonemia, Picozoa, and Katablephariodophyta (Table S5).

#### Supplementary Figures and Tables

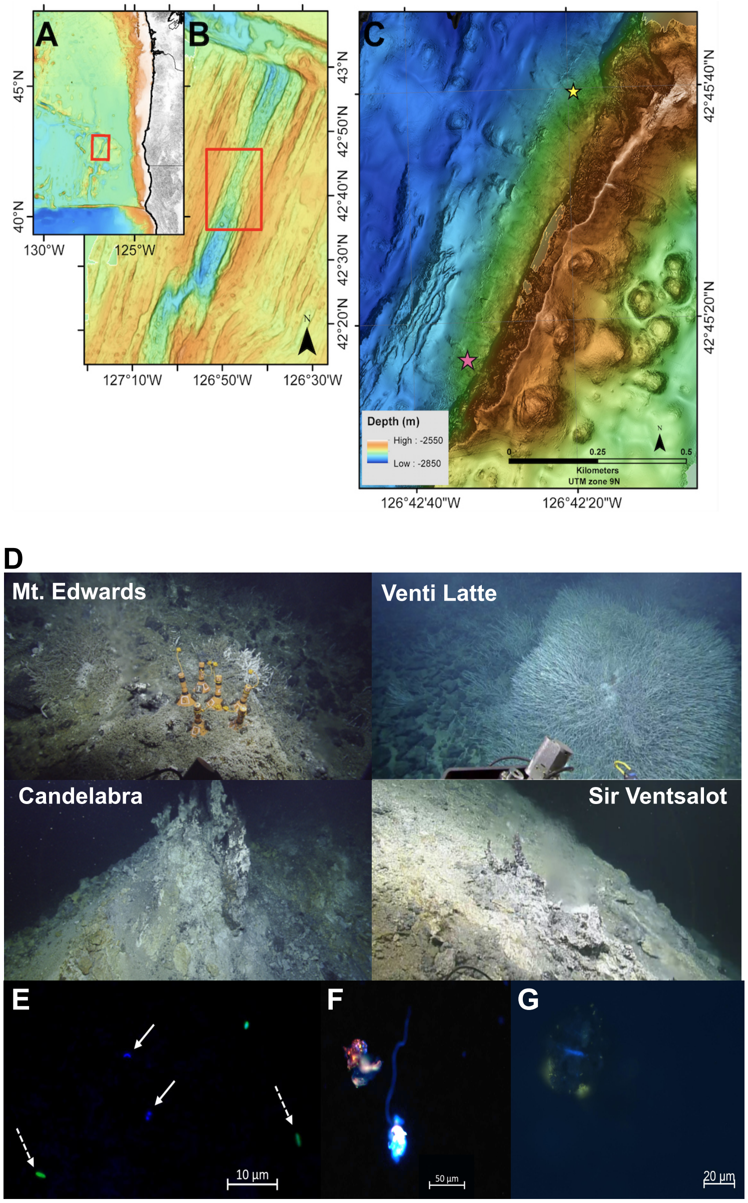


**Figure S1.** (**A-C**) Location of Sea Cliff and Apollo vent fields on the Gorda Ridge shown by the pink and yellow stars in **C**, respectively. Map modified from Clague *et al.* (21). (**D**) Images taken during sample collection from the four vent sites where grazing experiments were conducted. Copyright Ocean Exploration Trust, Inc. (**E**) Epifluorescence image from grazing experiment counts where solid arrows denote *in situ* bacteria and dashed arrows indicate fluorescently-labelled prey (FLP). The image demonstrates that the FLP were within the size range of the *in situ* microbial population. (**F-G**) Images taken of eukaryotic cells from the grazing experiments. Preserved sample material was set aside and filtered onto 0.8 μm filters to image vent associated protists under epifluorescence.

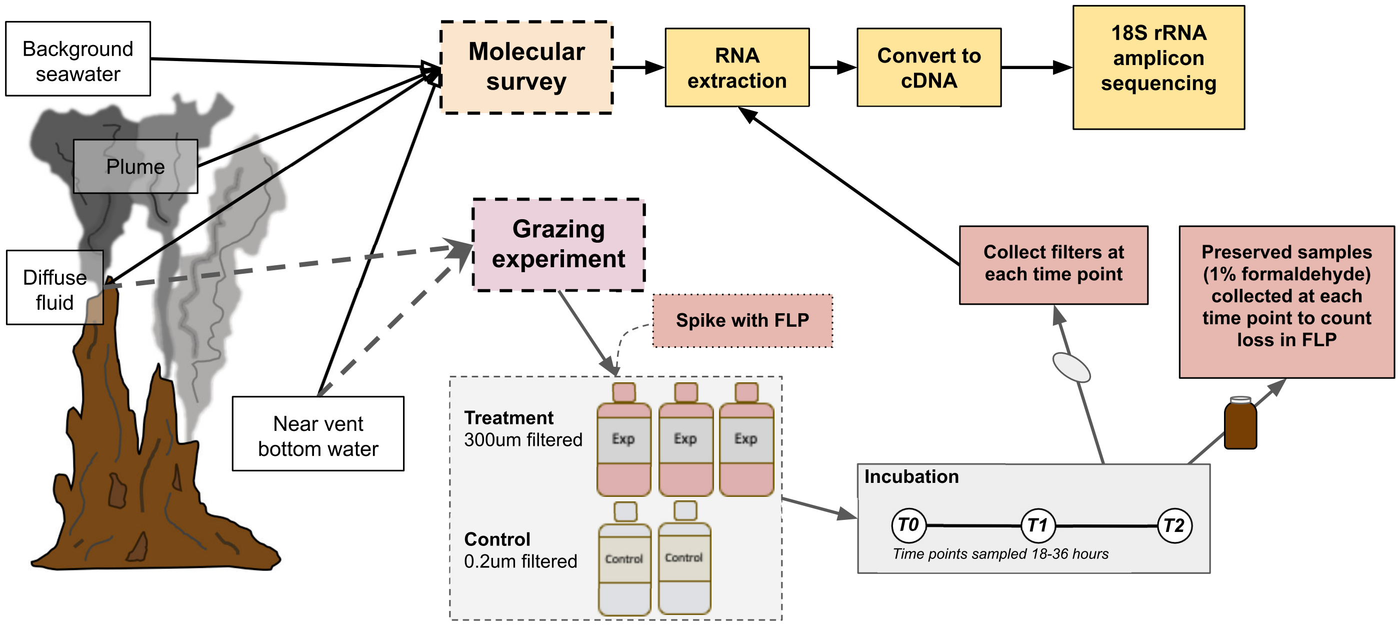


**Figure S2.** Schematic showing sample origin and general processing steps for molecular survey and grazing experiments. ROV Hercules collected *in situ* filters and fluid from diffuse fluid, plume (a couple of meters above the venting fluid), near vent bottom water, and background seawater.

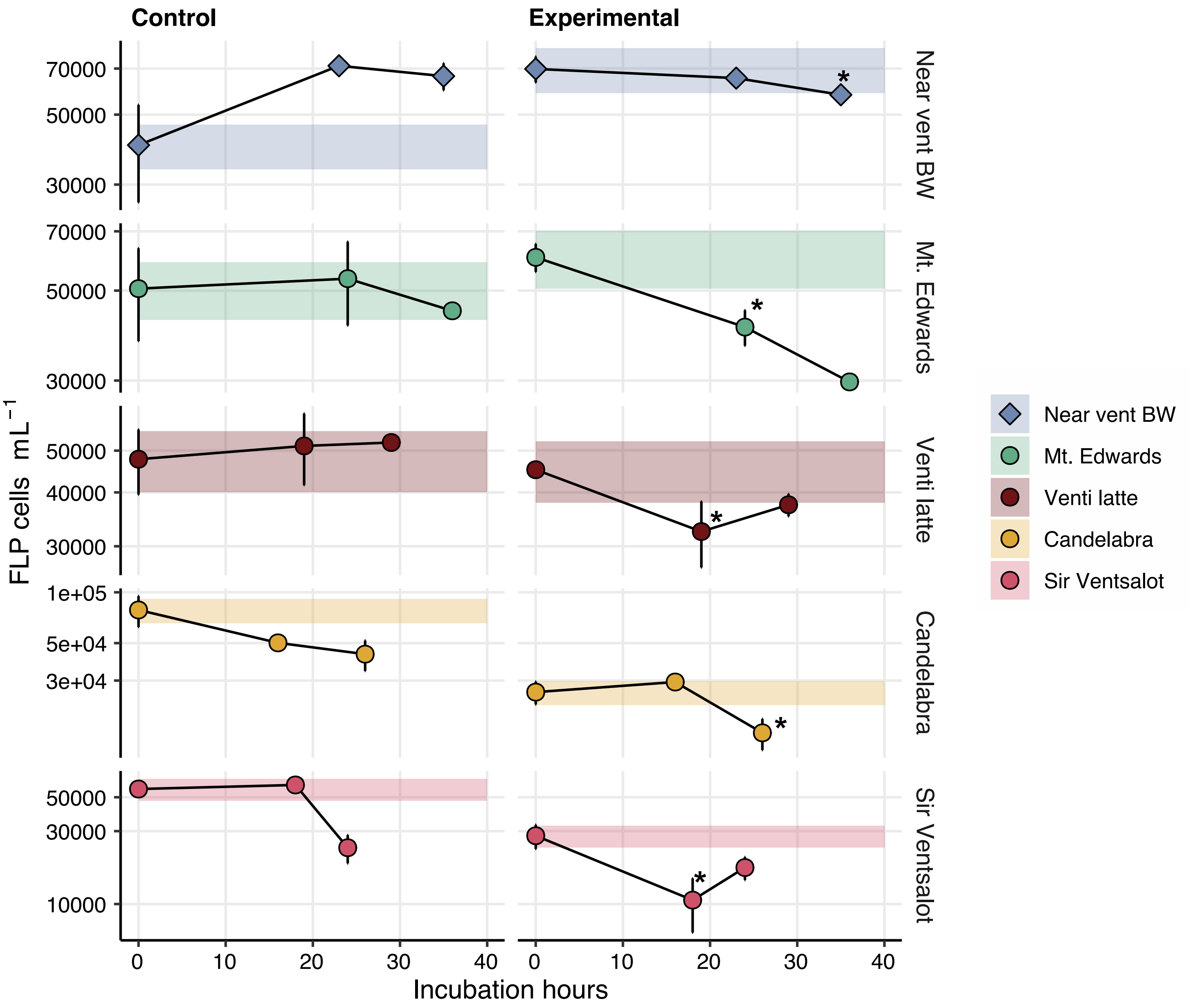


**Figure S3.** Loss in fluorescently-labeled prey (FLP) over time for each grazing experiment. FLP loss is shown for both the controls (left panel) and experimental treatments (right panel). Error bars represent the standard mean error and data points represent the average of replicate samples. Shaded area represents the determined microscopy error percentage above and below the $T_0$ time point; this metric serves to demonstrate significant changes in FLP concentration. FLP change in the control experiments (right) demonstrates that introduced FLPs did not disappear for other reasons besides grazing; for most experiments, FLP concentration in the control samples remained within the margin of error at the determined $T_F$ time point (left). Experiments where FLP loss at $T_1$ or $T_2$ fell below the microscopy error were used to calculate the extent and rate of grazing; labeled with an asterisk (*). See Materials and Methods for additional explanation.

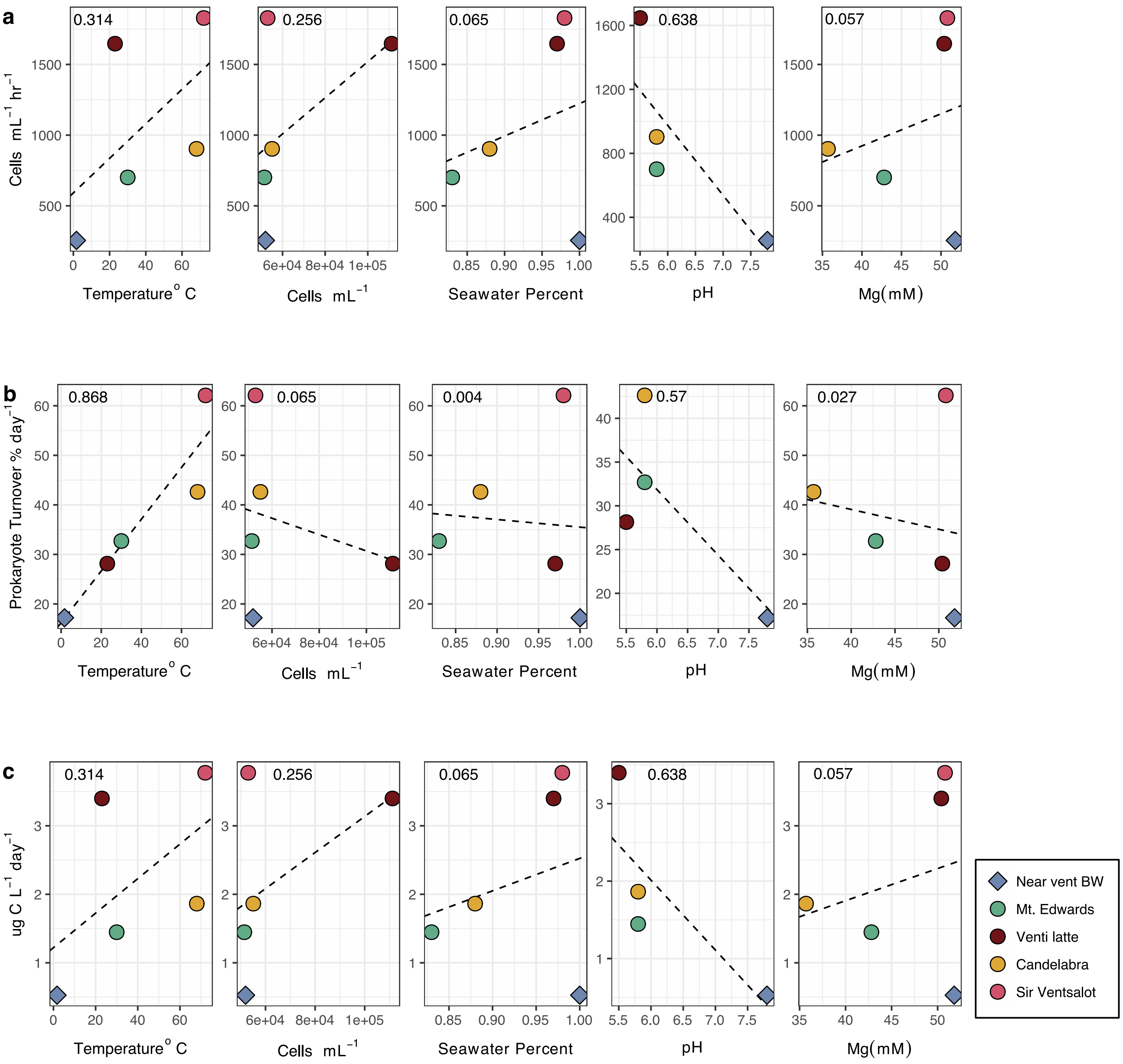


**Figure S4.** Relationship between measurements of protistan grazing pressure (top to bottom) and environmental parameters (from left to right): temperature, prokaryote concentration percent seawater of diffuse fluid, pH, and magnesium. Environmental parameters (x-axis) are shown in relation to (**a**) grazing rate in cells ml$^{-1}$ hour$^{-1}$, (**b**) prokaryote turnover percentage day$^{-1}$, and (**c**) µg C L$^{-1}$ day$^{-1}$ consumed. Dashed lines represent slope from linear regressions and r$^2$ values are listed in the top left corner of each plot. Environmental data were obtained from SUPR bag samples that were used for grazing experiments (Table 1). Estimates of grazing pressure (grazing rate, rate of prokaryote turnover, and rate of carbon consumed) do not strongly correlate with environmental parameters, with the exception of prokaryote turnover percent day$^{-1}$ with temperature.

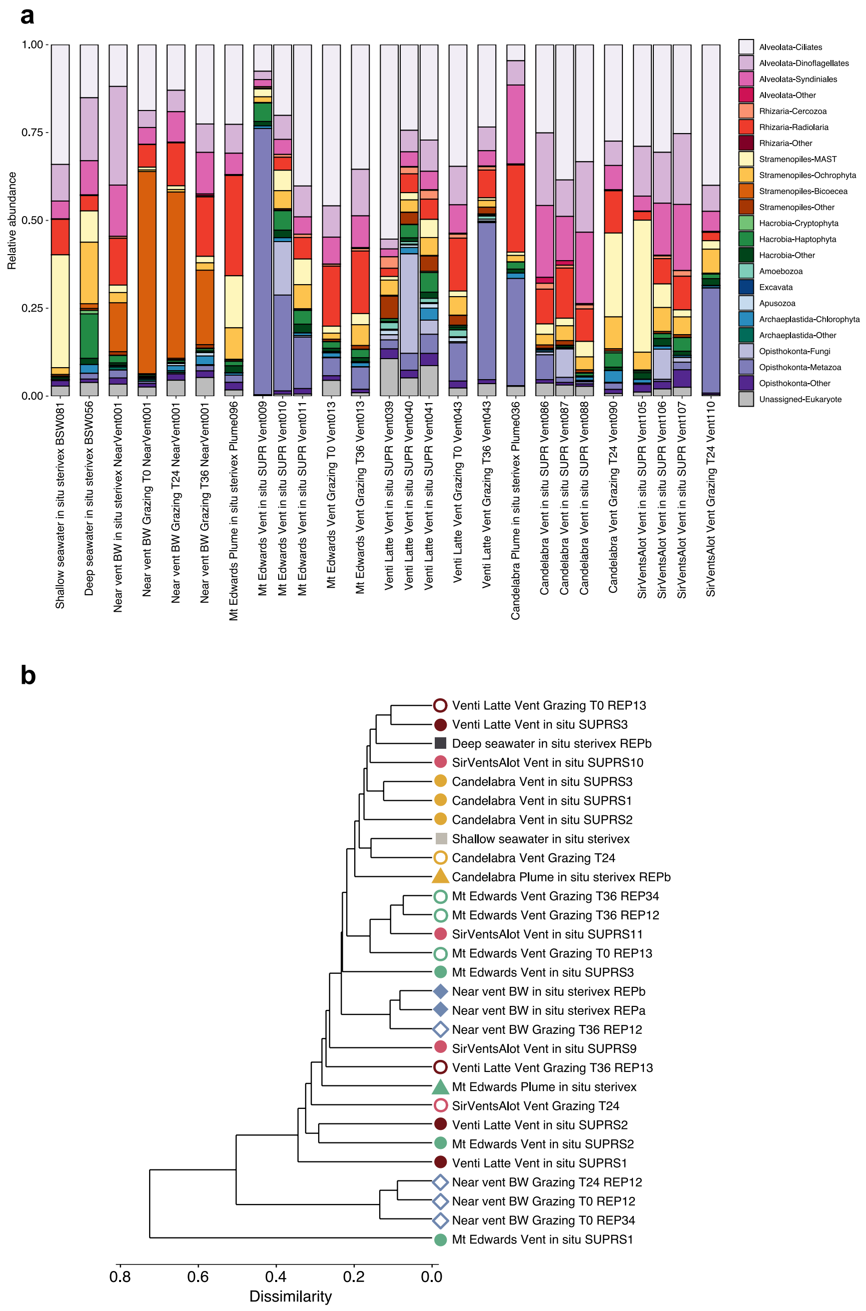


**Figure S5**. (**a**) Taxonomic breakdown of samples from the Gorda Ridge collected from background seawater, diffuse hydrothermal fluids, and associated grazing incubation experiments. Bar plot shows the relative sequence abundance for each sample (including replicates) and colors designate major protistan taxonomic groups, which has been manually curated (see *Materials and Methods*). Bar plot also varies from the main text by showing the relative sequence abundance of opisthokonta and unassigned sequences. (**b**) Average hierarchical clustering of all samples following relative abundance transformation and dissimilarity calculation. Colors and symbols are consistent with Figure 2.

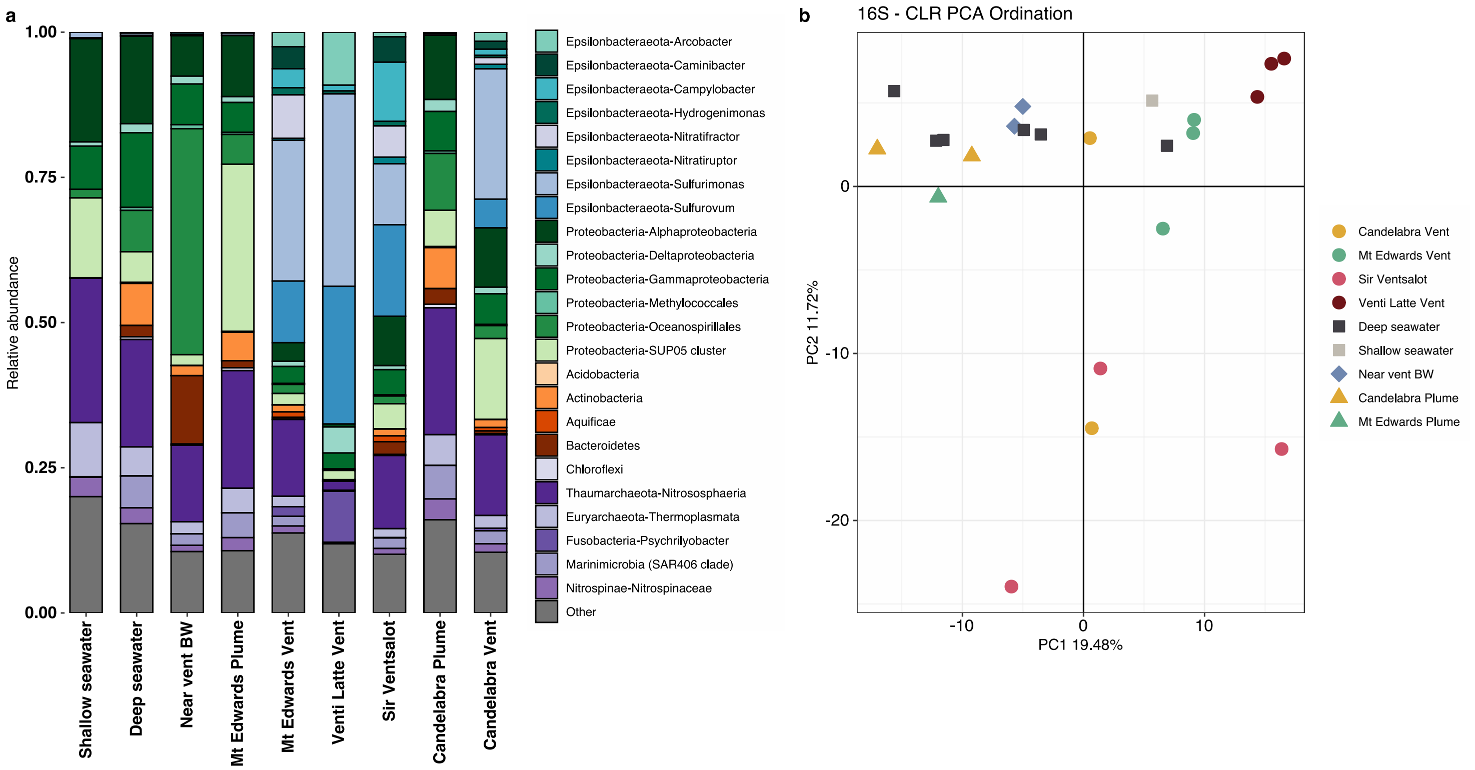


**Figure S6**. (**a**) Taxonomic breakdown of 16S rRNA gene amplicon results from Sea Cliff and Apollo hydrothermal vent fields derived from background and *in situ* vent samples. Colors denote bacteria or archaea taxonomic groups. Most samples are the result of averaging among 2 or 3 samples taken at the same time. The two leftmost samples originated from background shallow (150 m) and deep seawater (>2000 m). The "Other" category represents 16S rRNA gene-derived ASVs that were less than 0.1% in abundance or were manually removed due to known extraction kit representatives. (**b**) Ordination analysis of all samples, including replicates, from the 16S rRNA gene-derived sequence data. Data was center log-ratio transformed ahead of PCA analysis, similar to Figure 2b for the 18S rRNA-based ordination analysis.

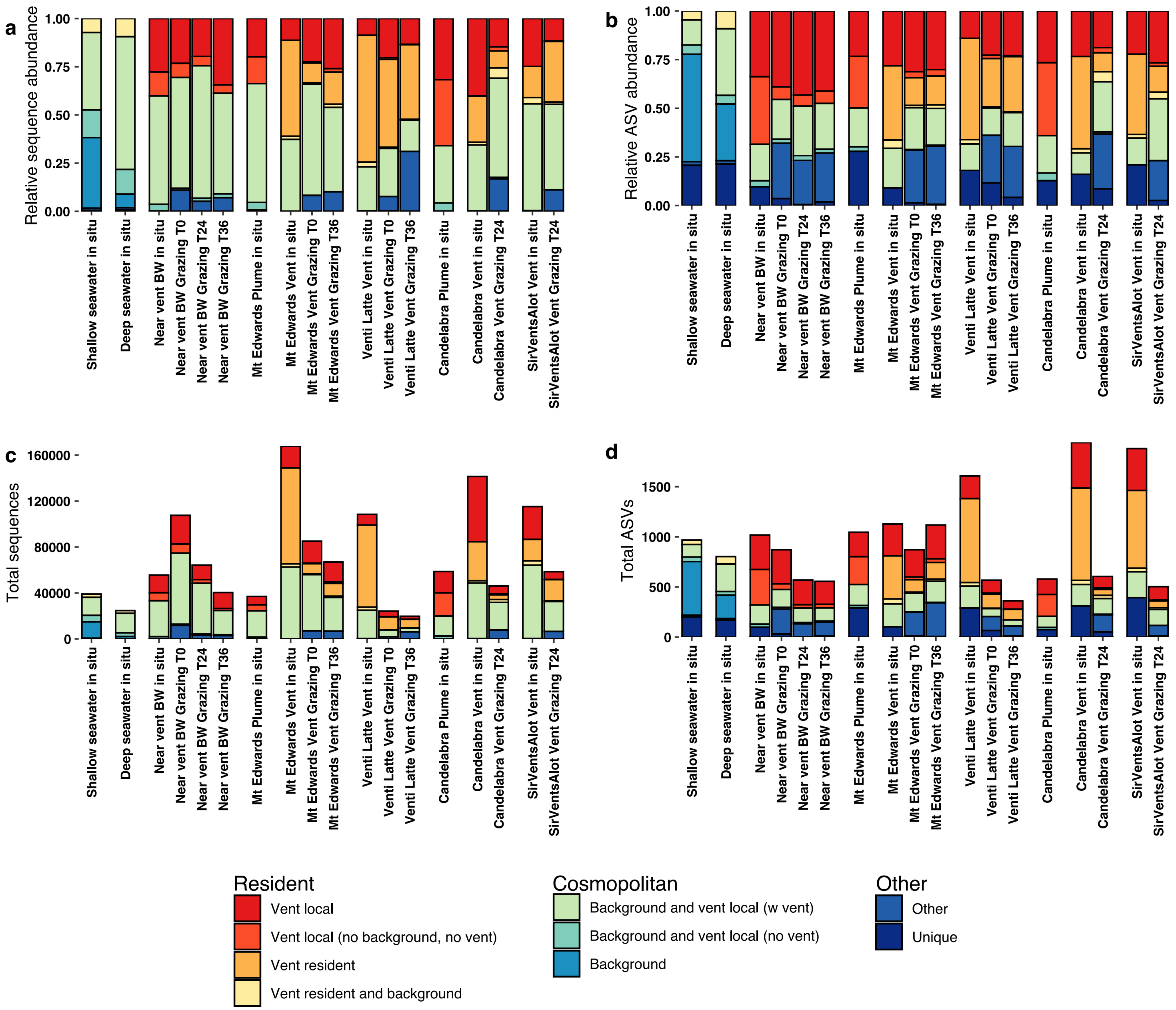


**Figure S7.** Distribution of protistan ASVs among background and vent sites (see main text for description of cosmopolitan versus resident). Here, the resident and cosmopolitan populations are further grouped by the presence of ASVs throughout the Gorda Ridge. Relative abundance of (**a**) sequences and (**b**) ASVs based on the distribution of ASV occurrence (denoted by color). Total number of (**c**) sequences and (**d**) ASVs based on the distribution of ASV occurrence.

***See Supplementary files for Tables S1-S6***

**Table S1.** Complete experiment details for each grazing incubation. Sample information (orange) rows list dive IDs and identifiers designated from EV Nautilus. Grazing incubation details (green) list start times and sampling time points (two per experiment) for each experiment and temperature of incubations. Due to the novel nature of these experiments, incubations were run at a variety of times, but were sampled at either approximately $T_{18}$ and $T_{24}$ or $T_{24}$ and $T_{36}$. Finally, the bottom columns (blue) list which time point was found to have a significant difference from the $T_0$ (based on microscopy error; see *Materials and Methods* and Figure S3), the fluorescently-labeled prey (FLP) cell concentration at $T_0$ and $T_F$, the average *in situ* prokaryote cell concentration, calculated mortality factor (m), grazing rate, and estimated prokaryote turnover percentage, results from carbon conversion estimates, and associated statistics.

**Table S2.** Sample names, metadata, and SRA IDs for all sequence samples in this study. Both 18S and 16S rRNA amplicon sequencing was conducted for this study, all sequences are submitted under SRA BioProject PRJNA637089.

**Table S3.** Total number of ASVs (top table) and sequences (bottom table) for each major protistan group. Columns list each sample type and represent the average across replicates and the sum across samples from the same grazing incubation.

**Table S4.** Total number of ASVs (top table) and sequences (bottom table) for each protist group at the class or family level. Taxonomic levels were curated to the class or family level as shown in Figure 3. Columns list each sample type and represent the average across replicates and the sum across samples from the same grazing incubation.

**Table S5.** The 10 most abundant ASVs within each protistan taxonomic group. Feature.ID reports the unique ASV identification, Distribution indicates if the ASV was found to belong to the resident or cosmopolitan population, and other columns report the taxonomic classification. The ASV size is the total number of sequences associated with the ASV.

**Table S6.** Summary of the significant 18S-16S ASV correlations derived from SPIEC-EASI results. The first four columns report the ASV Feature.ID and complete taxonomic name for the 18S and 16S ASV involved in the putative interaction. The following columns report the 18S ASV distribution and broader taxonomic classifications for both the 18S and 16S results. Finally, the weight reports the correlation value between the inferred relationships, the value reflects the strength of the interaction.
